# Raman microspectroscopy reveals unsaturation heterogeneity at the lipid droplet level and validates an *in vitro* model of bone marrow adipocyte subtypes

**DOI:** 10.1101/2022.10.03.510634

**Authors:** Josefine Tratwal, Guillaume Falgayrac, Alexandrine During, Nicolas Bertheaume, Charles Bataclan, Daniel N. Tavakol, Vasco Campos, Ludovic Duponchel, George Q. Daley, Guillaume Penel, Christophe Chauveau, Olaia Naveiras

## Abstract

Bone marrow adipocytes (BMAds) constitute the most abundant stromal component of adult human bone marrow. Two subtypes of BMAds have been described, the more labile regulated adipocytes (rBMAds) and the more stable constitutive adipocytes (cBMAds), which develop earlier and are more resilient to environmental and metabolic disruptions. *In vivo*, rBMAds are enriched in saturated fatty acids, contain smaller lipid droplets (LDs) and more readily provide hematopoietic support than their cBMAd counterparts. Mouse models have been used for BMAds research, but isolation of primary BMAds presents many challenges, and thus *in vitro* models remain the current standard to study nuances of adipocyte differentiation. No *in vitro* model has yet been described for the study of rBMAds/cBMAds.

Here, we present an *in vitro* model of BM adipogenesis with differential rBMAd and cBMAd-like characteristics. We used OP9 BM stromal cells derived from a (C57BL/6xC3H)F2-op/op mouse, which have been extensively characterized as feeder layer for hematopoiesis research. We observed similar canonical adipogenesis transcriptional signatures for spontaneously-differentiated (sOP9) and induced (iOP9) cultures, while fatty acid composition and desaturase expression of *Scd1* and *Fads2* differed at the population level. To resolve differences at the single adipocyte level we tested Raman microspectroscopy and show it constitutes a high-resolution method for studying adipogenesis *in vitro* in a label-free manner, with resolution to individual LDs. We found sOP9 adipocytes have lower unsaturation ratios, smaller LDs and higher hematopoietic support than iOP9 adipocytes, thus functionally resembling rBMAds, while iOP9 more closely resembled cBMAds. Validation in human primary samples confirmed a higher unsaturation ratio for lipids extracted from stable cBMAd-rich sites (femoral head upon hip-replacement surgery) versus labile rBMAds (iliac crest after chemotherapy). As a result, the 16:1/16:0 fatty acid unsaturation ratio, which was already shown to discriminate BMAd subtypes in rabbit and rat marrow, was validated to discriminate cBMAds from rBMAd in both the OP9 model *in vitro* system and in human samples. We expect our model will be useful for cBMAd and rBMAd studies, particularly where isolation of primary BMAds is a limiting step.

## 1 Introduction

Similar patterns of bone marrow adipocyte (BMAd) formation occur in vertebrate species (including rodents, rabbits, and humans). Specifically, a decreasing rate of BMAd has been observed relative to the decreasing size and lifespan of the animal, with larger skeletons containing more BM adipose tissue (BMAT) which extends farther into the skeleton (1). The distal parts of the skeleton (ie. peripheral bones in human, the distal tibia, paws, and caudal vertebrae in mice) predominantly contain adipocytic marrow interspersed with some hematopoietic cells, comprising the more stable constitutive BMAds (cBMAds) that appear just around birth. The proximal locations of the skeleton (ie. thoracic vertebrae in human and mouse, as well as proximal tibia and other long bones in mouse) contain predominantly hematopoietic marrow interspersed with the more labile regulated BMAds (rBMAds) (1,2, 36). The rBMAds are smaller in size and respond readily to induction of bone marrow adipogenesis through nutritional challenge (high fat diet or caloric restriction) or hematopoietic failure (irradiation or chemotherapy), or BMAd mass reduction in the context of cold exposuror increased hematological demand (e.g. phenylhydrazine). It is argued that cBMAds on the other hand, do not respond readily to environmental demands (1,3–6). Stromal cells and BMAds are tightly linked to BM hematopoiesis. Various stromal populations in mouse and human have shown to promote hematopoietic support while more mature BMAds correlate with reduced hematopoietic proliferation ((7,8) and reviewed in (9,10)). Meanwhile *Adiponectin* expressing cells within the BMAd lineage are beneficial to hematopoietic regeneration in most bones but not the caudal vertebrae of mice (11,12). These studies indicate the probability for population- and location-specific distinctions of BMAd maturation that specifically affect hematopoiesis and respond to hematopoietic demand (reviewed in (13)).

Isolation of BM stroma for differentiation and analyses of BMAds *in vitro* to study these processes in mouse models is challenging due to the limited amount of BMAd material obtained from each bone or bone segment. Moreover, this is compounded by the fact that the defining surface markers that would aid in purification of BMAd progenitor populations in homeostatic mice do not discriminate between the labile and stable BMAds (7). With such constraints, a promising alternative to the use of primary cells for *in vitro* studies are multipotent BM-derived stromal cell lines differentiated *in vitro*, especially when studies are possible at the single cell level to address heterogeneity (14). For this purpose, we developed a high-throughput image platform to monitor adipocytic differentiation though LD accumulation quantified by Digital Holographic Microscopy in live cells, which requires minimal handling to avoid perturbation of the culture (15). While not limited by end-point analysis or the introduction of handling or preparation biases, this technique does not provide information on the molecular composition of the cells. Methods capable of combining manipulation-free, label-free, single-cell imaging with analysis of molecular composition are thus of great interest to the field of adipogenesis at large. Raman microspectroscopy is a non-invasive and label-free method that does not require specific sample preparation, with a sub-cellular resolution on the scale of ~*μ*m that provides the molecular composition with spatial data, where standard methods provide bulk information (e.g. mass spectroscopic and chromatographic methods) (16). We thus hypothesized that combining lipid profiling by Raman microspectroscopy with the inherently heterogenous *in vitro* stromal adipocytic differentiation model would serve as a powerful tool to aid in the understanding of BM adipogenesis.

Specifically, LD formation (17) begins with fatty acids (FAs) that enter the cell via fatty acid transport proteins or fatty acid translocase (18,19), or are synthesized through endogenous de novo lipogenesis (DNL). FAs then enter a bioactive pool to form fatty acyl-CoA to be used by glycerolipid synthesis enzymes in the endoplasmic reticulum (ER) to form neutral lipids such as triacylglycerols (TAGs) and eventually coalesce and form LDs (20). If attached to the ER, LDs may grow in size through diffusion of newly synthesized lipids to the LD, and if unattached, through local synthesis or fusion of smaller LDs to larger ones. LDs increase in size during adipocytic differentiation and perilipins (PLINs) bind to promote their stabilization (21). Lipases mobilize neutral lipids in LDs for metabolic energy by fatty acid oxidation, which is triggered through nutritional, hormonal, or inflammatory activation. Adipose triglyceride lipase (ATGL) catalyzes the initial step of intracellular TAG hydrolysis followed by hormone sensitive lipase (HSL) and monoacylglycerol lipase (MGL) into glycerol moieties (22,23). FAs are necessary for energy production and lipid synthesis for cellular signaling and membrane formation. Despite their importance, increased concentrations of non-esterified FAs can be detrimental contributing to lipotoxicity, and thus TAG hydrolysis and FA cycling are carefully regulated (24,25). As illustrated in Figure S1, the primary product of DNL is palmitic acid (16:0), in the family of saturated FAs that have cytotoxic effects and induce reactive oxygen species (ROS) (26). The monounsaturated FA (MUFA) oleic acid (18:1) may counteract this effect possibly by activating esterification of palmitic acid into TAGs and lipid droplet storage (27). Palmitic acid accumulation is therefore prevented by increased desaturation to palmitoleic acid (16:1n-7) or elongation to stearic acid (18:0) and further desaturation to oleic acid (28). Notably, oleic acid inhibits palmitic-acid dependent osteoclastogenesis and palmitic acid is found to be increased in BM serum of osteoporotic women (29). In fact, BMAT expansion throughout the red marrow with age and disease is associated with bone loss and fracture risk, while cBMAd formation is positively associated with bone accrual during early development through correlative and descriptive studies (30).

While concrete definitions and BMAd classification markers are being established (30), the oldest standard for classifying the two known types of BMAds is by the composition of their lipid content (31, 32). The first reports by Mehdi Tavassoli showed that the adipocytic-rich yellow marrow -now denominated cBMAd-from the os calcis of rabbits was higher in the proportion of unsaturated FAs (palmitoleic −16:1n-7- and oleic −18:1n-9-acids), whereas fatty acid composition of BMAds from the adipocyte-poor red marrow of thoracic vertebrae (rBMAds) was richer in saturated FAs (palmitic −16:0- and stearic −18:0-acids) (33). In recent years, primary rat cBMAds from tail vertebrae and distal tibia were likewise shown to contain a higher proportion of unsaturated fatty acids in caudal vertebrae or distal tibia as compared to rat rBMAds isolated from either lumbar vertebrae or femur plus proximal tibia combined (1). Congruently MRI, ^1^H-MRS, and gas chromatographic data from human subjects consistently showed that BM adipose tissue (BMAT) from sites of red marrow contained highly saturated lipids, compared to the mostly unsaturated lipids detected at yellow marrow sites (1, 33–35). From murine samples, it is considerably more challenging to obtain sufficient BMAT for analysis due to the size of the animal as well as the amount of BMAT present in the bones (32). To our knowledge, the lipid composition of murine BMAds has not been described to date. However, between BMAT of red and yellow marrow there is a conservation of the differences in both the 16:1/16:0 and 18:0/18:1 fatty acid unsaturation ratios relative to total lipid content across species (summarized and compiled with our data in Table 1 and Figure S1).

**Table 1:** Fatty acid unsaturation ratios in rat, rabbit, human, and OP9 cells from red and yellow BMAds. The 16:1/16:0 and 18:1/18:0 unsaturation ratios were calculated from the proportion of total fatty acids. Rat BM data from Scheller et al. 2015 Supplementary Data 1 (regulated: femur and proximal tibia, constitutive: distal tibia). Rabbit BM data from Tavassoli 1977 Table 1 (regulated: vertebrae, constitutive: os calcis). Human original data from Figure 7 (regulated: iliac crest post-chemotherapy, constitutive: femoral head) and OP9 cell original data from Figure 6 (regulated: spontaneous, constitutive: induced) from HPLC analysis. Additional measurements in human BM have been made by MRI spectroscopy but cannot be compiled with these data due to the different methodology (35, 64). c: constitutive, r: regulated.

| Origin | BMA<sub>d</sub> type | 16:1/16:0 | c/r | 18:1/18:0 | c/r | Reference |
| --- | --- | --- | --- | --- | --- | --- |
| *Rat* | constitutive regulated | 0.25<br>0.09 | 2.8 | 1.18<br>0.81 | 1.5 | Scheller et al. 2015 |
| *Rabbit* | constitutive regulated | 0.53<br>0.16 | 3.3 | 7.52<br>2.68 | 2.8 | Tavassoli et al. 1977 |
| *Human* | constitutive regulated | 0.19<br>0.11 | 1.7 | 7.91<br>4.40 | 1.8 | original data |
| *OP9 cells* | induced spontaneous | 0.84<br>0.24 | 1.6 | 4.74<br>2.33 | 2.0 | original data |

**Table 2:**
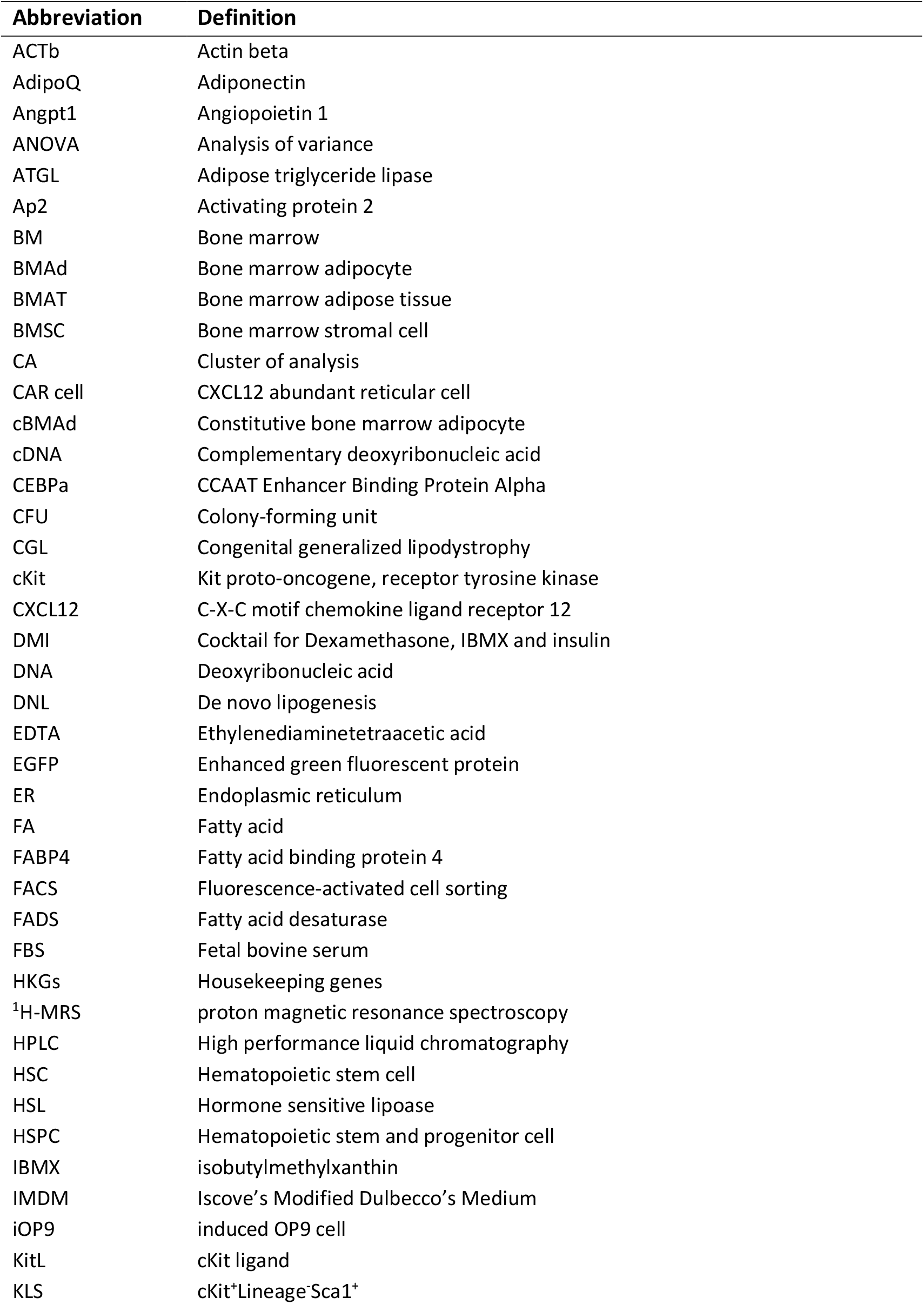
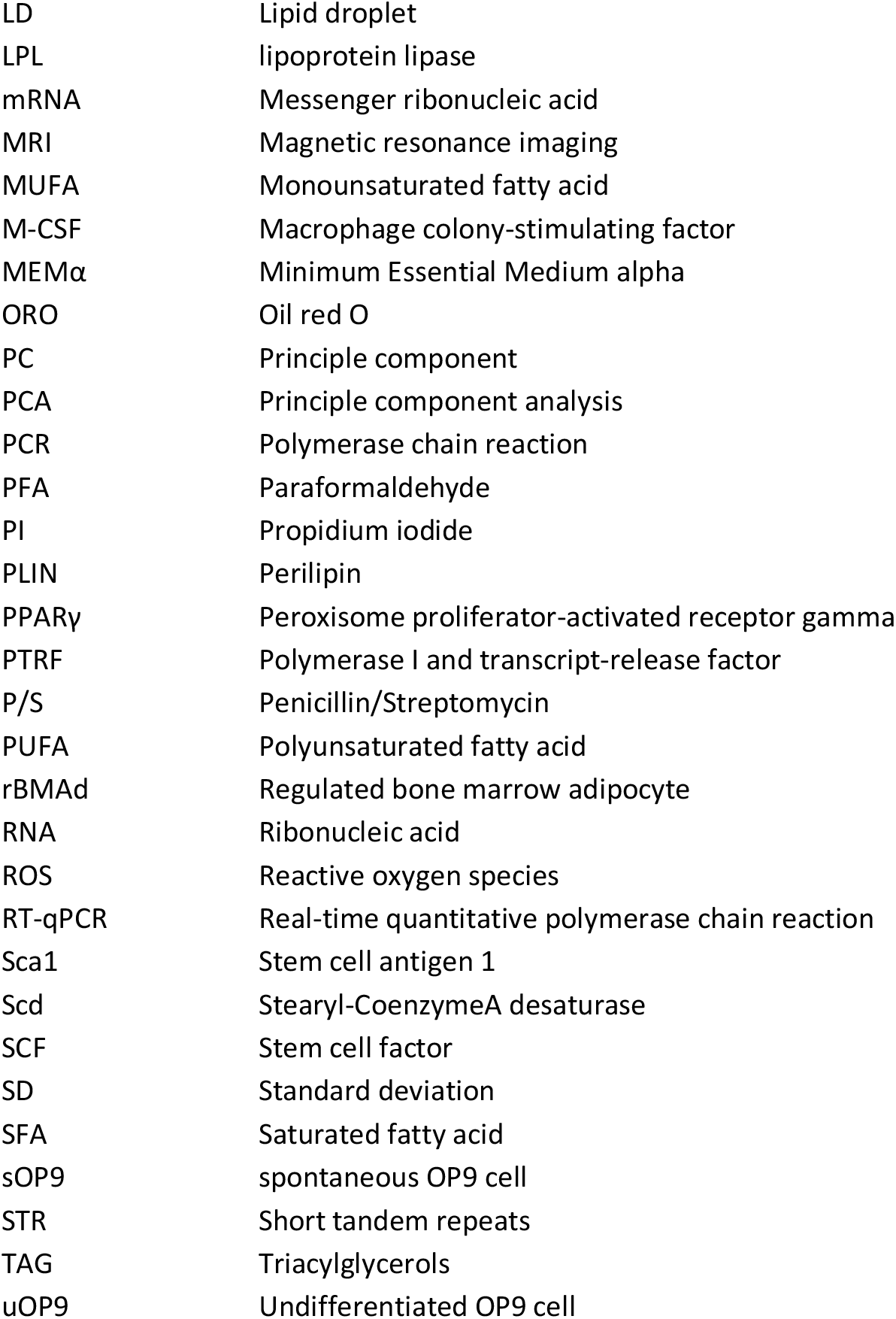
List of general abbreviations.

**Table 3:** List of fatty acid abbreviations.

| Abbreviation | Definition | Formula | IUPAC Name |
| --- | --- | --- | --- |
| 14:1 | Myristoleic acid | C<sub>14</sub>H<sub>26</sub>O<sub>2</sub> | (Z)-tetradec-9-enoic acid |
| 16:0 | Palmitic acid | C<sub>16</sub>H<sub>32</sub>O<sub>2</sub> | hexadecanoic acid |
| 16:1n-7 | Palmitoleic acid | C<sub>16</sub>H<sub>30</sub>O<sub>2</sub> | (Z)-hexadec-9-enoic acid |
| 16:1n-10 | Sapienic acid | C<sub>16</sub>H<sub>30</sub>O<sub>2</sub> | (Z)-hexadec-6-enoic acid |
| 18:0 | Stearic acid | C<sub>18</sub>H<sub>36</sub>O<sub>2</sub> | octadecanoic acid |
| 18:1n-9 | Oleic acid | C<sub>18</sub>H<sub>34</sub>O<sub>2</sub> | (Z)-octadec-9-enoic acid |
| 18:2n-6 | Linoleic acid | C<sub>18</sub>H<sub>32</sub>O<sub>2</sub> | (9Z,12Z)-octadeca-9,12-dienoic acid |
| 20:4n-6 | Arachidonic acid | C<sub>20</sub>H<sub>32</sub>O<sub>2</sub> | (5Z,8Z,11Z,14Z)-icosa-5,8,11,14-tetraenoic acid |

In this study, we harness the inherent adipogenic potential of murine BM-derived OP9 stromal cells to further dissect intrinsic properties of BMAds using standard techniques for lipid profiling at the cell population-level, and Raman microspectroscopy for lipid profiling at the single-adipocyte level. We subjected OP9 cells to basal culture conditions or to a standard adipogenic cocktail, allowing the OP9 cells to differentiate spontaneously though serum exposure and confluency (sOP9) or through specific induction of the adipogenic differentiation program (iOP9). To test the hematopoietic supportive capacity of OP9 cells from these conditions, we assessed their interaction with a hematopoietic component in short-term co-culture assays with hematopoietic stem and progenitor cells (HSPCs). We show that a full biochemical induction of the adipogenic differentiation program induces iOP9s to more closely resemble cBMAds in their lipid content, lipid droplet size and hematopoietic support than the sOP9 adipocytes that develop upon simple serum induction, which retain rBMAd-like properties. Furthermore, we validated the significance of our unsaturation ratio-driven cBMAd versus rBMAd definition. For this, we compared for the first time primary human samples from iliac crest samples drawn after hematological aplasia, which represent the most extreme case of rBMAd remodeling (36), to femoral specimens from hip replacement surgery, which represent one of the best characterized cBMAd depots (37, 38).

## 2 Methods

### 2.1 Cell culture

All stromal cells (MS5, C3H10T1/2, MC3T3-E1-4, MC3T3-E4-24 bone marrow stromal lines, AFT024 and BFC012 stromal lines from fetal liver (39) or embryonic stem cell-derived fibroblasts as in (40, 41) were cultured in complete medium consisting of Minimum Essential Media alpha (MEMα) with GlutaMa™ (Gibco, catalog no. 32561) and 1% Penicillin/Streptomycin (P/S, Gibco, catalog no. 15140) supplemented 10% fetal bovine serum (FBS, Gibco, catalog no. 10270-106) at 37°C and 5% CO_2_ with media changed every 2-3 days, and differentiated as described below (42). For subsequent analysisOP9 cells were plated in 24-well plates (Falcon) with CaF_2_ substrates (Crystran) for Raman microspectroscopy and RT-qPCR, in 6-well plates (Falcon) for HPLC analysis, and in flat-bottom tissue culture-treated 96-well plates (Falcon) for co-culture assays. OP9 cells were plated subconfluently at a density of 5,000 cells per cm^2^ (undifferentiated OP9 cells) or plated at confluency of 20,000 cells per cm^2^ in complete medium (spontaneous OP9 cells) for either 7 or 17 days, as specified in figure legends. Both OP9 cells and OP9-EGFP cells (14) were authenticated by the ATCC on June 24, 2021, by short tandem repeat (STR) analysis, indicating an exact match of the ATCC reference OP9 cell line (CRL-2749). OP9 cells are available from the ATCC (CRL-2749), but this source has not been tested for adipogenesis or hematopoietic support in our laboratory.

#### 2.1.1 *In vitro* differentiation and quantification

All stromal lines were plated at 20,000 per cm^2^ and cultured to confluency in 1% gelatin-coated plates (prepared overnight at 4°C), then allowed to spontaneously differentiate over time by confluency or induced toward adipocytic or osteogenic differentiation via a standard differentiation cocktail as in (14, 42). The adipogenic induction cocktail consisted of complete medium supplemented 1*μ*M dexamethasone (Sigma, catalog no. D4902), 5*μ*g/ml insulin (Sigma, catalog no. I0516), and 0.5mM isobutyl-methylxanthine (IBMX, Sigma, catalog no. I5879). After four days, the adipogenic induction medium was changed to a maintenance medium consisting of complete medium with insulin and dexamethasone only. Media was changed every 3-4 days with aliquots prepared fresh from stock solutions (IBMX in DMSO, insulin in PBS, dexamethasone in ethanol) kept at −20°C in the dark. Short-term differentiation experiments were performed at day seven of adipocytic differentiation with minimal culture manipulation as in (15), and long-term experiments were carried out on day 17. Neutral lipids were stained at day 17 of culture with Oil Red O for quantification of adipogenesis in 96-well plates. First, cells were gently washed with phosphate buffered saline (PBS, Gibco, catalog no. 10010015) then fixed for 10min at room temperature with 4% paraformaldehyde (PFA) diluted in PBS from 32% stock solution, and gently washed three times with PBS. 100*μ*l filtered Oil Red O solution freshly prepared from stock solution (Sigma, catalog no. 01391-250ML) diluted 3:2 with distilled water was added to the wells. Cells were incubated with Oil Red O at room temperature for 45min on a shaker, then washed gently with PBS three times and light transmission micrographs were obtained. 100*μ*l isopropanol was then added to the wells and incubated at room temperature. After 10min, 70*μ*l of the solution was transferred to a new 96 well plate and OD measurements read at 520nm with isopropanol as background. Osteogenic differentiation was performed by addition of dexamethasone 1*μ*M, 2-phospho-L-ascorbic acid 50*μ*g/ml (1000x stock in PBS), glycerophosphate 10mM (100x stock in PBS) and 1,25-hydroxyvitamine D3 0.01*μ*M (stock 1000x in ethanol) for 28 days. Every 3-4 days differentiation media was changed. All media changes were made with fresh stock aliquots. Efficiency of osteoblastic differentiation was determined with Alizarin Red stain (Sigma) in stromal cells fixed as described above for O Red Oil stains. Alizarin Red staining solution (alizarin red 2% in distilled water, filtered through 45-*μ*m pore, pH adjusted to 4.0-4.3 with NH4OH and filtered again) was then added for 10-15 minutes until precipitates were visible. Alkaline Phosphatase development kits were purchased from Promega (S3771), and OP9 stains were performed according to manufacturer’s instructions in lightly fixed cells (4%PFA for 1 minute at room temperature) by dilution of NBT in AP staining buffer (100mM Tris HCl pH9.5, 50mM MgCl2, 100mM NaCl and 0.1% Tween-20). Pictures were taken immediately after staining.

### 2.2 Raman microspectroscopy

Fixed 24-well plates were kept in PBS at 4°C and shipped to the University of Lille. Raman acquisitions were done on a LabRAM HR800 equipped with an immersion objective (Nikon, obj x100, numerical aperture = 1, Japan) and a diode laser λ=785 nm. The laser power at the sample was 30mW. The lateral resolution was 1-2*μ*m. Spectral acquisition was made in the 400–1800cm^-1^ range and spectral resolution was 4cm^-1^. The acquisition time was set at 60s per spectrum. Raman spectra were processed using Labspec software (HORIBA, Jobin-Yvon, France). The water immersion objective focused the laser on the center of individual lipid droplets where one spectrum corresponds to one adipocyte lipid droplet. In total 2944 spectra were measured over 120 sOP9-adipocytes (on average 24 spectra per adipocyte) and 2971 spectra over 138 iOP9-adipocytes (with an average of 21 spectra per adipocyte). The number of spectra per well was between 60 to 110. The unsaturation ratio was calculated as ratio of area under the curve of bands 1654cm^-1^ / 1441cm^-1^. The optical image of the adipocyte was saved for each acquisition. The diameter of LDs was evaluated from the optical image by using the software FIJI (43).

The instrument is also equipped with a XYZ motorized stage which allows the acquisition of Raman images. Raman images were acquired using the point-by-point imaging mode (Pt-Img). The laser beam was focused perpendicular to the sample surface. Acquisition time was set to 3sec (×2) for each spectrum. The laser beam was stepped in two dimensions (x and y), and a spectrum was recorded at each position (x,y). The step was set to 1 *μ*m between 2 positions. The Pt-Img mode generated x×y spectra.

### 2.3 High-performance liquid chromatography

Cells were fixed with PFA as described for the adipocytic quantification above. After washing, 500*μ*l of PBS was left in the wells, plates were sealed with parafilm and stored at 4°C until processing for HPLC analysis. Cells were detached from the plate with 1mL (2 × 0.5ml) of PBS and lipids extracted with chloroform/methanol (2:1; v/v) under agitation for 30min at room temperature. The resulting lipid extract was then subjected to a saponification, followed by a fatty acid derivatization into naphthacyl esters as described previously (44). Fatty acid derivatives were applied into the HPLC Alliance system (2695 Separations Module, Waters, Saint-Quentin-en-Yvelines, France) equipped of an autosampler (200ml-loop sample), a photodiode array detector (model 2998), and the Empower software for data analyses. Fatty acid derivatives were eluted on a reverse phase YMC PRO C18 column (3mm, 4.6×150mm, 120Å) by using two solvent systems: a) methanol/acetonitrile/water (64:24:12; v/v/v) and, b) methanol/dichloromethane/water (65:28:7; v/v/v) under identical conditions reported previously (44). Fatty acid derivatives were detected at 246 nm and quantified by an external standard curve realized with the naphthacyl 19:0 derivative. Note that 19:0 was also used as internal standard to evaluate FA recoveries. Lipidomics LC-HRMS standards and solvents are provided in Table S1.

### 2.4 Real-time quantitative PCR

At day seven or 17 of adipocytic differentiation, RNA was extracted with Trizol and cDNA was synthesized using Taq DNA polymerase (catalog no. 10342020, Life Technologies) according to manufacturer’s instructions. Quantitative PCR was performed in technical triplicates using the Fast SYBR Green qPCR Mastermix with 250 *μ*M of primer concentration (Applied Biosystems) on QuantStudio 6 (Life Technologies). Primers were pre-validated for high efficiency and internally tested not to vary upon OP9 differentiation. The geometric mean of housekeeping genes *(RPL13* and *YWHAZ)* was used as reference to calculate the fold expression of each gene relative to the samples of undifferentiated OP9 cells as described for the delta-delta normalization method (45, 46). Primer sequences obtained from Microsynth are listed in Supplementary Table S2.

### 2.5 Bone marrow extraction and HSPC sorting

Total bone marrow was extracted from eight-week-old B6 ACTb-EGFP females housed in 12-hour day-night light cycles and provided ad-libitum sterile food and water as described previously, in accordance to Swiss law and with the approval of cantonal authorities (Service Veterinaire de l’Etat de Vaud) and complying with ARRIVE guidelines. Total bone marrow cells were extracted from femur, tibia and pelvis by crushing using a mortar and pestle in ice-cold PBS supplemented 1mM ethylenediaminetetraacetic acid (EDTA, catalog no. 15575020, Thermo Fisher Scientific). The samples were dissociated and filtered through a 70*μ*M cell strainer (catalog no. 352350, Falcon), lysed for 30sec at room temperature in red blood cell lysis buffer (Biolegend, catalog no. 420301), washed with ice-cold PBS-EDTA and centrifuged at 1300rpm for 10min at 4°C. For HSPC sorting prior to co-culture with OP9 stroma, the cell pellet was stained with 50*μ*l biotinylated ‘lineage’ antibody cocktail (BD, catalog no. 558451) in 1ml PBS-EDTA per six bones for 15min on ice. The sample was washed with PBS-EDTA and stained with 50*μ*l magnetic beads of the same kit in PBS-EDTA for 10min on ice. The samples were then washed and filtered through a 70*μ*m cell strainer (catalog no. 352350, Falcon) prior to lineage depletion using the AutoMACS Pro (Miltenyi Biotec, USA). After depletion the negative fraction was resuspended in an antibody mix containing antibodies against Streptavidin-TxRed (1:200), cKit-PECy7 (1:200), Sca1-APC (1:100) and PI (1:1000). Cell sorting was performed on a FACSAria Fusion (Becton Dickinson, USA) cell sorter.

### 2.6 *In vitro* HSPC co-culture

OP9 cell seeding as detailed in Section 2.1, was followed by a short-term *in vitro* adipocyte differentiation of OP9 cells, and then a gently wash with PBS and pre-warmed Iscove’s Modified Dulbecco’s Medium (IMDM, catalog no. 12440053, Gibco) supplemented 10 % FBS and 1% Pen/Strep (P/S) was added. The washing was done using the Caliper Sciclone ALH 3000 (Caliper Life Sciences, USA) and consisted of four cycles of removing 140*μ*l media and adding 140**μ**l fresh IMDM media, to dilute the previous media so as to keep mature adipocyte detachment to a minimum. The wells were left with 100**μ**l IMDM 10% FBS and 1% P/S. HSPCs extracted from B6 ACTb-EGFP mice were plated at a ratio of 1:10 initial OP9 cells (in volume of 100**μ**l IMDM, totaling 200**μ**l volume per well).

After seven days of co-culture, the 96-well plates were removed from the incubator and placed on ice. To count non-adherent cells, the cells in suspension were retrieved through repeated washes and stained with CD45-PacBlue at final concentration of 1:200, PI at 1:1000 and 5**μ**l CountBright beads (catalog no. C36950, Invitrogen) directly added to the media. Without any further manipulation, the plates were analyzed using the High Throughput Sampler (HTS) module of a LSRII (Becton Dickinson, USA) flow cytometer. For accuracy, the settings were programmed to mix the wells thoroughly before sample uptake. To count adherent cells, a separate set of duplicate plates were taken from the incubator, the media manually removed, and wells washed once with 200*μ*l PBS. 40*μ*l of Trypsin EDTA (0.5 %, catalog no. 25300-054, Gibco) was added to each well and incubated for five minutes at 37°C. 160*μ*l of ice-cold PBS containing FBS (to neutralize the trypsin), CD45-APCCy7 (1:200), PI (1:1000) and 5*μ*l CountBright beads were then added to the plates. The plates were stained on ice and directly analyzed with flow cytomet.

#### 2.6.1 In vitro conditioned media preparation and culture

Conditioned media was prepared for four different conditions as follows. For the undifferentiated condition, undifferentiated OP9 cells were seeded in six-well plates at confluency (20,000 OP9 cells/cm^2^) in IMDM (catalog no. 12440053, Gibco) supplemented 10% FBS and 1% P/S (basal IMDM) and conditioned medium was harvested after two days in culture and stored at −20°C. For the spontaneous differentiation conditions, confluent OP9 cells were grown in MEMα as described for the short-term differentiation assays except that at day five media was changed to IMDM following three washes; conditioned medium was then harvested after two days and stored at −20°C. For the induced adipocytic conditions, confluent OP9 cells underwent the short (5 day) or long (17-day) DMI adipogenic induction protocols before changing media to basal IMDM for two additional days of culture; conditioned medium was then harvested and stored at −20°C. Subsequently 2000 live-sorted HSPCs were plated per well in 96-well round-bottom plates in the absence of stroma. Different ratios of conditioned-to-basal IMDM media were added. After two-day culture the cells were stained with Annexin V, Propidium Iodine (PI) and CD45 to select for live CD45^+^ hematopoietic cells via flow cytometric analysis with an LSRII cytometer (Becton Dickinson, USA). CountBright beads were added to count absolute cell numbers as described above.

#### 2.6.2 Cobblestone formation assay

OP9 cells were plated at 20,000 cells/cm^2^ in gelatin-pre-coated 6-well plates and cultured with α-MEM media supplemented with 10%FBS and 1%P/S. The cells were induced to differentiate using the standard differentiation cocktail DMI, which was changed two times per week. After seven days of differentiation, the media was washed three times with PBS and changed to basal IMDM. On the same day, 200 sorted KLS cells were added into each well. After 7 days of co-culture the hematopoietic colonies were manually counted (47) and scored the following way: (i) shiny colonies (defined by majority of round and bright cells, as they float above the OP9 stromal layer, with no more than 10 cobblestone cells visible per colony; (ii) cobblestone colonies (defined as darker ‘cobblestone’-like cells, because they are below the OP9 stromal layer) should not contain more than 20% of shiny cells; and (iii) mixed colonies that contain both shiny and cobblestone cells(48–50).

### 2.7 Clinical samples

This study complied with the Declaration of Helsinki and the local ethical authorities (CER-VD). All patients signed a specific consent for the reuse of biological samples in the context of our study. For HPLC, iliac crest BM aspirates from female and male patients undergoing treatment for acute myeloid leukemia (between ages 48 and 73, n=5) or surgical debris from patients undergoing hip replacement surgery (between ages 56 and 80, n=4) were received for analysis from the Lausanne University Hospital (Centre Hospitalier Universitaire Vaudois, CHUV). Floating adipose tissue, serum, and oil fractions were isolated and processed for lipid extraction and processed as described for OP9 lipidomics.

### 2.8 Statistical Analysis

Values are shown as mean plus or minus the standard deviation or standard error of the mean as indicated. Student’s t-test was performed for all experiments when comparing two conditions only, or a Two-Way ANOVA when comparing multiple conditions, with P-values indicated for statistical significance.

Raman spectra were processed by using the PLS Toolbox and the MIA toolbox (v8.7 eigenvector Research, Inc., USA). The spectra were baseline corrected, normalized and mean centered. Two processes were done on Raman spectra. First, cluster analysis (CA) was performed using Ward’s method. CA clustered individual Raman spectra into 3 categories (saturated-rich lipids, unsaturated-rich and mixture of saturated and unsaturated lipids). Second, principal component analysis (PCA) was used which highlighted the difference in spectral features between the spontaneous and induced conditions. All data are reported as mean ± standard deviation. All statistical analyses were performed using GraphPad Prism (version 8.0.0, GraphPad Software). Mean values were compared by Mann and Whitney test, or two-way analysis of variance (ANOVA) followed by Bonferroni’s post hoc test. Statistical significance was accepted for P<0.05, and reported.

## 3 Results

### 3.1 Differential adipocyte lipidation and similar transcriptional signature upon adipogenic induction by confluency (spontaneous) versus DMI cocktail (induced)

We chose murine BM-derived OP9 stromal cells as a model for this study because of their propensity to differentiate into adipocytes while displaying both hematopoietic support and dual adipogenic and osteogenic differentiation capacities as compared to other commonly used murine stromal lines including BM-derived MS5 and C3H10T1/2 lines or MC3T3 subclones (Figure S2A-C and (42)). Note retention of alkaline phosphatase activity in confluent and fully mature, lipidated OP9-derived O Red Oil positive adipocytes (Figure S2D), reminiscent of properties assigned to hematopoietic supportive AP+ reticular adventitial cells in the bone marrow (51) and possibly of AdipoCARs (52).

The non-clonal OP9 stromal cell line (53, 54) was originally derived from the calvaria of newborn osteopetrotic mice ((C57BL/6xC3H)F2-op/op) deficient in macrophage colony-stimulating factor (M-CSF), and were shown optimal for studies of hematopoietic cell development and differentiation sensitive to M-CSF (55, 56). OP9 cells have been described as preadipocytes with multilineage differentiation capacity which readily generate mature adipocytes, and express the transcriptional activators CCAAT/enhancer binding proteins (C/EBP) α and β, together with the master regulator of adipocyte differentiation peroxisome proliferator activated receptor-γ (PPARg), and perilipin (PLIN), a phosphoprotein associated with the surface of lipid droplets (LDs) (21, 54). Thanks to their multipotency, hematopoietic support, and adipocytic differentiation capacities, OP9 cells are thus ideal for *in vitro* studies on BM adipogenesis. Due to their non-clonal nature and inherent heterogeneity, they serve as a BM model for studying BMAd differentiation at the single cell level.

*In vitro*, the OP9 cell line spontaneously differentiates into adipocytes upon confluency in the presence of serum (spontaneous OP9 adipocytes, sOP9), a phenomenon that can be more robustly promoted by exposure to a classical adipocytic differentiation cocktail consisting of insulin, dexamethasone, and 3-isobutyl-1-methylxanthine (induced OP9 adipocytes, iOP9) (21, 53). After 17 days in culture, both sOP9 and iOP9 culture conditions contained numerous differentiated OP9-adipocytes (Figure 1) with a significantly greater prevalence of Oil Red O staining of lipids in the induced condition (iOP9) (p<0.01) as compared to control baseline undifferentiated OP9 cells plated at sub-confluency (uOP9) (Figure 1G). Spontaneous OP9-adipocytes formed under the minimal confluency condition (sOP9) and stained with Oil Red O at intermediate levels between undifferentiated uOP9 cells and induced iOP9 cultures (Figure 1G). No significant amount of mature adipocytes could be visualized in the uOP9 condition (Figure 1A,D). Adipocytes were defined as previously described, where cells containing at least four identifiable lipid droplets were accounted as mature adipocytes (57). Morphologically, both sOP9 and iOP9 adipocytes stained with Oil Red O at day 17, but iOP9 adipocytes constituted the majority of the cultures and often contained large lipid droplets, while sOP9 adipocytes were sparser and their lipid droplets seemed smaller.

**Figure 1:**
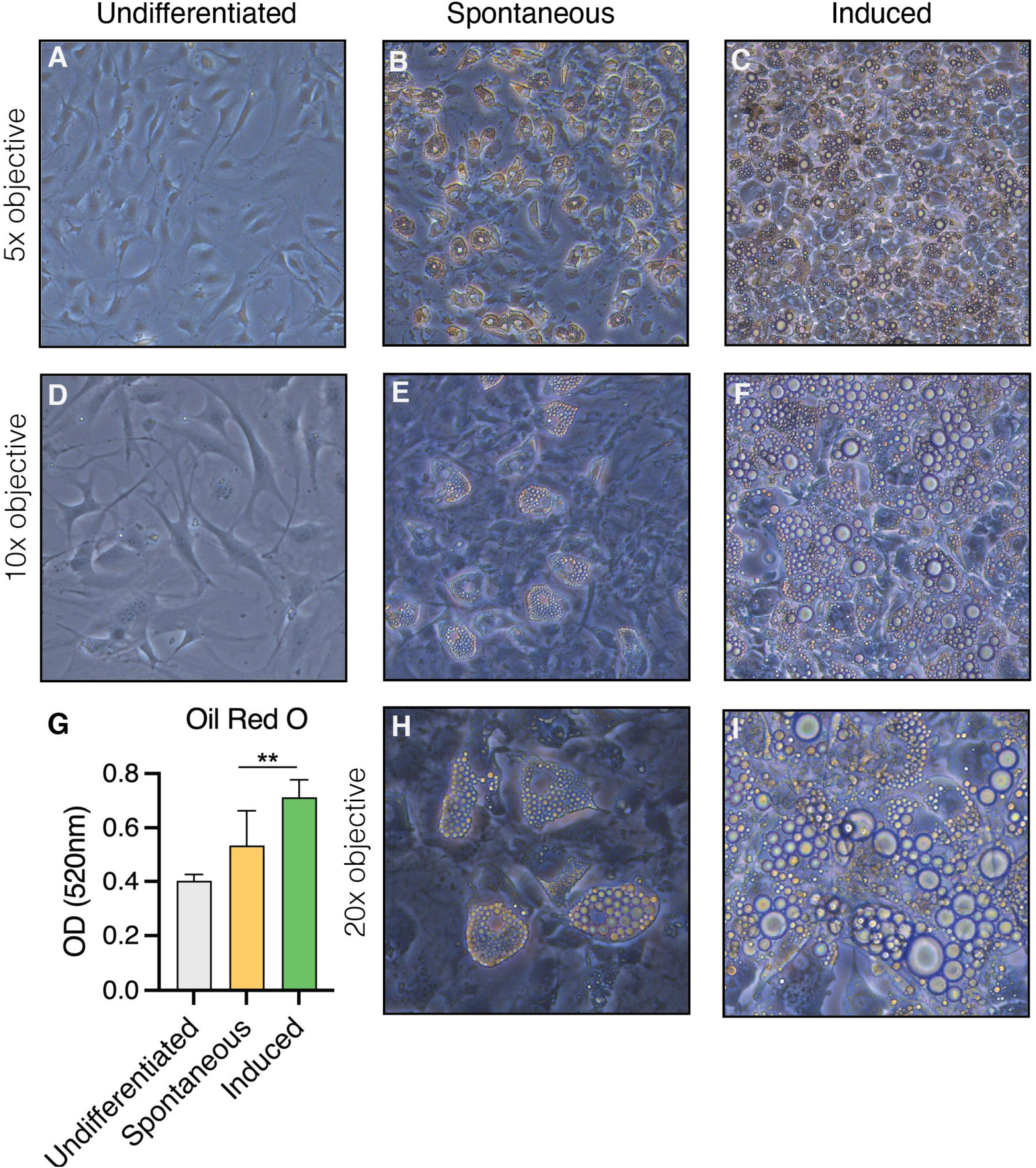
Heterogeneity in the formation of lipidated adipocytes upon induction of adipogenesis by confluency and serum exposure (spontaneous) versus DMI cocktail (induced). OP9 cells were (**A**, **D**) passaged subconfluently and remained undifferentiated on day 17 of culture, (**B**, **E**, **H**) differentiated spontaneously upon confluency in the presence of serum for 17 days, or (**E**, **F**, **I**) induced with a classical adipogenic differentiation cocktail in the same serum-containing conditions after confluent plating and 17 days in culture. (**G**) Oil red O measurements show the highest adipogenic differentiation in the induced condition at day 17 in culture. **P<0.01 by student’s t-test. Error bars represent mean ± s.d. (n=3 biological replicates). DMI: 1μM dexamethasone, 0.5mM isobutyl-methylxanthine, 5μg/ml insulin.

Given the intermediate phenotype of sOP9 adipocytic cultures, containing a mixture of mature adipocytes and non lipidated stromal cells likely including preadipocytes, we wished to determine whether the transcriptional profile and lipid composition of these cultures more resembled the undifferentiated uOP9 cultures or the highly lipidated iOP9 adipocytic cultures. We thus compared transcriptional adipocyte differentiation markers by RT-qPCR and the global fatty acid (FA) composition by high performance liquid chromatography (HPLC) for the three conditions.

RNA transcripts for the canonical adipocyte differentiation and maturation program *(Fabp4, Adipoq, Cebpa, Pparg, Lpl, Pnpla2* -also known as *Atgl-)* in the sOP9 condition most resembled iOP9 adipocytic cultures, with average gene expression more than 10 times higher for the sOP9 and iOP9 conditions than for the control uOP9 condition (Figure 2A). These results were further validated in the OP9-EGFP cell line used in previous studies (Figure S3A-B, (14)). Further analysis of the desaturation and elongation pathway for fatty acid synthesis showed a similar pattern (Figure 2B), with overall higher expression for the sOP9 and iOP9 adipocytic conditions as compared to the uOP9 undifferentiated condition. Of note RNA transcripts for *Sds1* and *Fads2* were however significantly between sOP9 and iOP9 cultures. We thus conclude that, at the population level, expression of genes related to the adipocyte differentiation and fatty acid desaturation programs is similar for both sOP9 and iOP9 adipocytic conditions after 7 days of culture, suggesting that the transcriptional program is triggered in sOP9 cells prior to lipid droplet accumulation.

**Figure 2:**
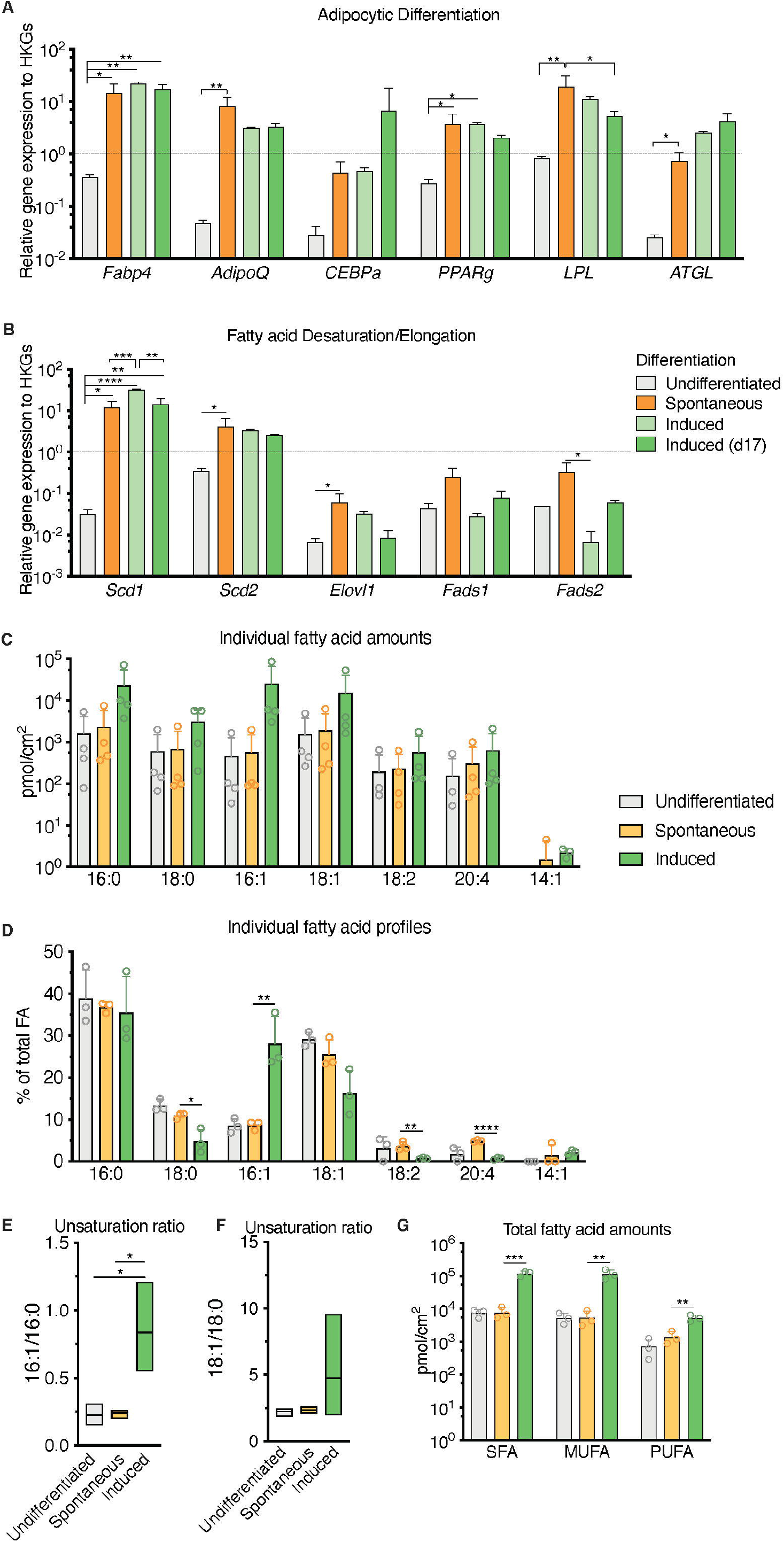
Population level differences in lipid composition are more marked than differences in the canonical adipocytic transcriptional signature for spontaneous versus induced cultures. (**A**) RT-qPCR markers of adipogenesis are increased in induced OP9-adipocytes. (**B**) Desaturase expression is highest in day 17 induced OP9-adipocytes while one of the elongases in fatty acid *de novo* synthesis is decreased. ****P<0.0001, ***P<0.001, **P<0.01, *P<0.05 by Bonferroni’s multiple comparisons test. Error bars represent mean ± s.d. (n=3 independent experiments). (**C**-**G**) HPLC analysis of spontaneously differentiated or induced OP9 adipocytes revealed (**C**) nuances of the most abundant saturated, mono-, and polyunsaturated lipid species, with (**D**) overall total higher unsaturation content in induced versus spontaneous conditions. (**E**) Percentage of individual fatty acid species reveals (**F**) a greater 16:1/16:0 unsaturation ratio in the induced condition. (**G**) Unsaturation ratios 18:1/18:0 were not significantly different. ****P<0.0001, ***P<0.001, **P<0.01, *P<0.05 by ANOVA test with Bonferroni’s multiple comparisons adjustment. Error bars represent mean ± s.d. (n=3 independent experiments). HKGs: housekeeping genes.

Conversely, total FA content and profiles were similar for control uOP9 cells and the sOP9 adipocytic condition as measured by HPLC analysis. Classical adipogenic induction of iOP9 cells resulted in a 16-fold increase of total FA content as compared with sOP9 cells or uOP9 cells (2.4×10^5^±2.2×10^4^pmol/cm^2^ versus 1.5×10^4^±7.0×10^3^pmol/cm^2^ and 17.4×10^4^±3.7×10^4^pmol/cm^2^ respectively, Figure 2C). FA profiles revealed a preferential enrichment of MUFAs in iOP9 cells (47% MUFAs versus 38% in uOP9s and 36% in sOP9 cells), mostly to the detriment of PUFAs (only 2% in iOP9 cells versus 6% PUFAs in uOP9s and 10% in sOP9 cells, p<0.0001) (Figure 2D, G). In terms of individual FAs, the most striking finding was the marked high ratio of palmitoleic acid (16:1) in iOP9 cells, representing 29% of total FA (versus 9% in uOP9 or sOP9 cells, p=0.007) (Figure 2D, E). This result was associated to a reduction of 18 carbon FAs in iOP9 cells and a trend towards a consistent reciprocal enrichment of 18 carbon FAs in sOP9 cells: stearic acid (18:0, sOP9 11% vs. iOP9 4%, p=0.02), oleic acid (18:1, sOP9 26% vs. iOP9 16%, p=0.07) and linoleic acid (18:2 n-6, sOP9 4% vs. iOP9 1%, p=0.007). The most abundant PUFA species was arachidonic acid (20:4 n-6). In relative terms, linoleic (18:2 n-6) and arachidonic acid (20:4 n-6) were significantly enriched in the sOP9-adipocytes at 5% of total FA while making up only 1% of total FA in iOP9-adipocytes. Notably, the ratio of palmitoleic acid (16:1) versus its saturated form (stearic acid, 16:0) in iOP9 adipocytes was three times higher than in sOP9 adipocyte cultures or uOP9 control cells (Figure 2E). The ratio of oleic acid (18:1) versus its saturated form (stearic acid, 18:0) showed a similar but non-significant trend (Figure 2F). These results are very similar to those obtained on rats’ tibiae when comparing lipid profiles from constitutive adipocytes versus regulated adipocytes (1). Our data is also congruent with the results shown for rabbit BMAds when comparing FA composition of rabbit BMAs from rBMAd-rich versus cBMAd-rich sites ((33) and summarized in Table 1).

Overall, these results indicate that, at the population level, sOP9 adipocytic cultures share transcriptional similarities in the adipocyte differentiation program with iOP9 cells, but are less lipidated and present lower abundance of unsaturated FAs. This constellation would be consistent with our hypothesis that sOP9 adipocytic cultures present similarities with regulated BMAds, and thus an enrichment in saturated FA, as compared to cBMAd-like iOP9 adipocytic cultures. Given the heterogeneous lipidation of sOP9 adipocytic cultures and the inevitably bulk nature of our RT-PCR and HPLC-based analysis, we set out to validate our hypothesis through a single cell analysis pipeline that could directly compare sOP9 and iOP9 adipocytes on their relative FA acid composition at the single cell level.

### 3.2 Raman microspectroscopy detects unsaturated spectra at the single lipid droplet level in spontaneous and induced OP9-derived adipocytes

Raman microspectroscopy represents a powerful yet non-invasive and label-free method to assess the lipid composition of adipocytes *in vitro*. The unsaturation ratio is recognized as a measure of the proportion of unsaturated FA obtained by the ratio of the peak area assigned to unsaturated bonds (C=C) divided by the peak area assigned to saturated bonds (CH2) (Equation 1) (58–61).

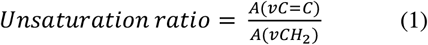

**Equation 1:** Lipid unsaturation ratio measured by Raman microspectroscopy.

Our feasibility test with Raman microspectroscopy imaging showed that it was possible to acquire Raman spectra at single lipid droplet (LD) resolution in our model (Figure 3). After 17 days of culture, Raman spectra, representative of the molecular composition of individual LDs, were acquired for sOP9 and iOP9 adipocytes through high resolution Raman imaging. No analysis was performed for the undifferentiated (uOP9) condition as it presented with very few, if any, LDs. Raman images of individual adipocytes were processed by PCA. The PCA shows that a mixture of spectra exists between the adipocytes but also within the LDs themselves, as illustrated by the Raman images (Figures 3B, 3E). In Figure 3C, the PC1 scores (96.68%) show a representative spectrum rich in unsaturated lipids for the iOP9 adipocyte shown in Figure 3A-B. In Figure 3F, the PC1 scores (95.69%) show a spectrum representative of a saturated rich lipids for the sOP9 adipocyte shown in Figure 3D-E. Figure S4 shows the additional variation captured by PC2 and PC3 (<5%) for the same two adipocytes shown in Figure 3. We conclude that LDs composed of predominantly unsaturated-rich or saturated-rich lipids can be identified through Raman microspectroscopy within OP9-derived adipocytes with single LD resolution.

**Figure 3:**
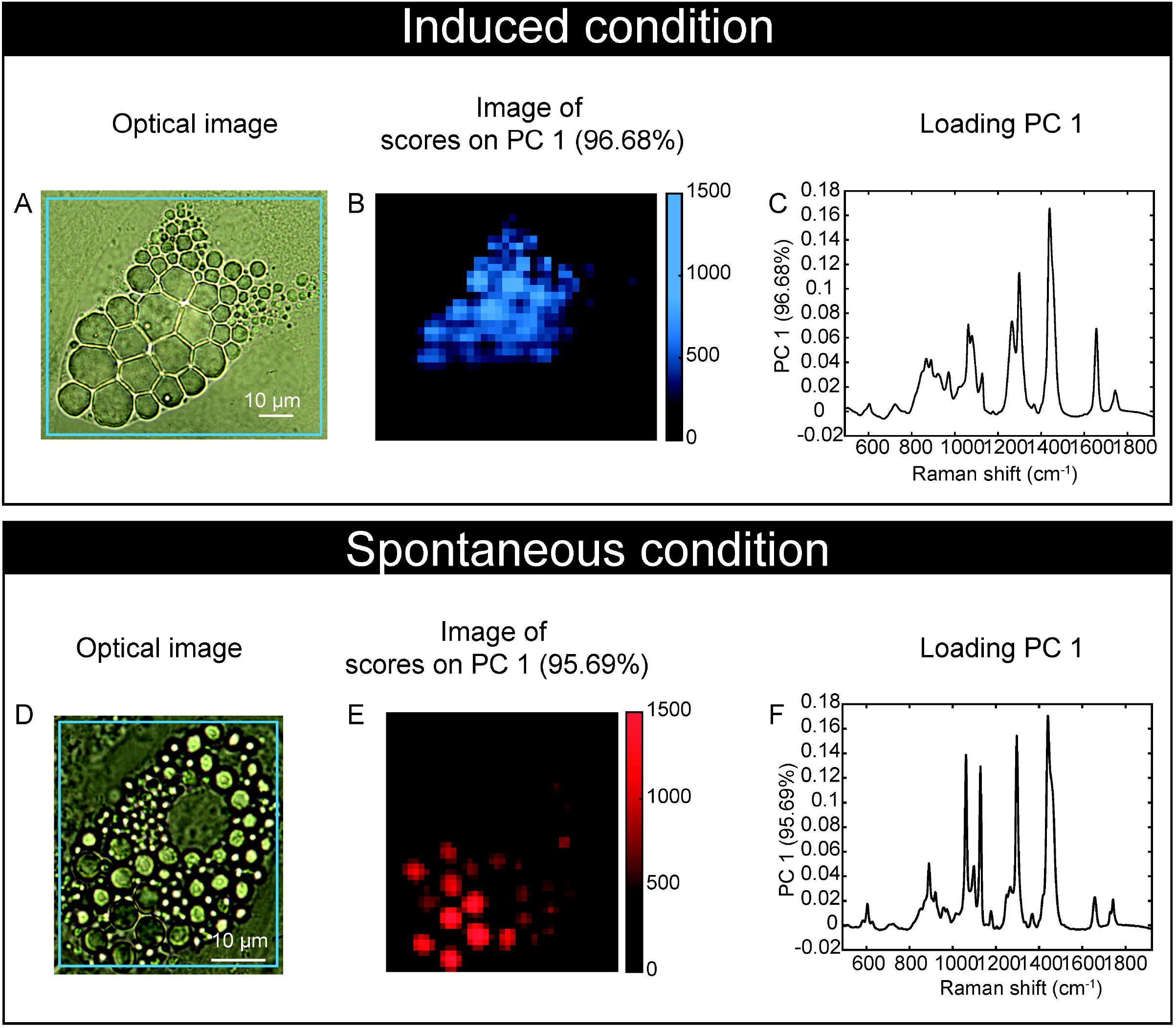
Raman microspectroscopy detects unsaturated spectra at the single lipid droplet level in spontaneous and induced adipocytes. (**A**) An optical image of an adipocyte in the induced condition (iOP9) analyzed by Raman imaging. (**B**) Raman image of the scores of PC1 in induced condition. Pixels in black represent a score of PC1 equal to zero, which corresponds to the absence of PC1. Pixels in shades of blue correspond to the contribution of PC1. (**C**) Loading PC1 is characteristic of a spectrum of unsaturated lipid-rich and captures more than 95% of the variation. (**D**) An optical image of an adipocyte in spontaneous condition (sOP9) analyzed by Raman imaging. (**E**) Raman image of the scores of PC1 in spontaneous condition. Pixels in black represent a score of PC1 equal to zero, which corresponds to the absence of PC1. Pixels in shades of red correspond to the contribution of PC1. (**F**) Loading PC1 is characteristic of a spectrum of saturated lipid-rich. The Raman images of the scores and the loadings of PC2 and PC3 capture less than 2% and are presented in supplementary files (Figure S4).

### 3.3 Raman microspectroscopy reveals differential predominance of unsaturated spectra in large lipid droplets from induced adipocytes

In order to determine if significant differences exist in the unsaturation ratio of FAs accumulated within sOP9 versus iOP9 adipocytes, we thus proceeded to the acquisition of Raman spectra from individual LD in both conditions and throughout 3 separate experimental campaigns (iOP9: n = 138 adipocytes and 2971 spectra; sOP9: n = 120 adipocytes and 2944 spectra; lateral resolution 1-2*μ*m), and combined all data for aggregated principal component (PC) analysis in both conditions. Figure S5 shows the classification of spectra using unsupervised hierarchical cluster analysis, which predicted three main categories: saturated-rich, unsaturated-rich and mixture. The representative spectrum of each category is shown in Figure 4A. PCA was then performed on the averaged spectra per adipocyte. The PC1 loading plot contains characteristic spectra where the positive peaks correspond to saturated lipids (1061, 1129, and 1296 cm^-1^, represented in red for Figures 4A-B) and the negative peaks correspond to unsaturated lipids (1080, 1267, and 1655 cm^-1^, represented in blue for Figures 4A-B) (58). PC1 captures 78.20% of the variability of the overall dataset, and separates the Raman spectra based on the saturated-rich and unsaturated-rich profiles (Figure 4B). On the PCA score plot, each point represents an averaged spectrum per adipocyte (Figure 4C). Most of the spectra from iOP9-adipocytes (81%) form a cluster in the negative score along PC1, while the spectra of sOP9-adipocytes are scattered along PC1. This indicates that the molecular composition of LDs in sOP9-adipocytes is heterogeneous compared to iOP9-adipocytes, and that iOP9-adipocytes are enriched in unsaturated-rich lipids. Score plots in Figure 4C and 4D are equivalent, but the points in Figure 4C are colored according to PC1 scores, and the points in Figure 4D are colored according to the culture condition. Specifically, the points in dark red and dark blue in Figure 4C correspond to Raman spectra of saturated-rich and unsaturated-rich lipids, while intermediate colors point to Raman spectra of a rather balanced mixture of saturated and unsaturated lipids. PCA analysis shows that LDs in iOP9-adipocytes are mainly rich in unsaturated lipids, while LDs in sOP9-adipocytes are composed of a rather balanced mixture of unsaturated and saturated lipids (Figure 4D). However, when comparing the averaged raw unsaturation ratio (based on averaged spectra per adipocyte) we found no significant difference between iOP9 adipocytes and sOP9 adipocytes (Figure 4E). At the adipocyte level, both conditions have a similar averaged unsaturation ratio. This result is explained by PCA analysis which shows that LDs of both conditions are not solely composed of either uniformly saturated or unsaturated lipids, but rather by different mixtures thereof. We thus took advantage of the 1-2*μ* resolution of our setup and performed the analysis at the single LD level.

**Figure 4:**
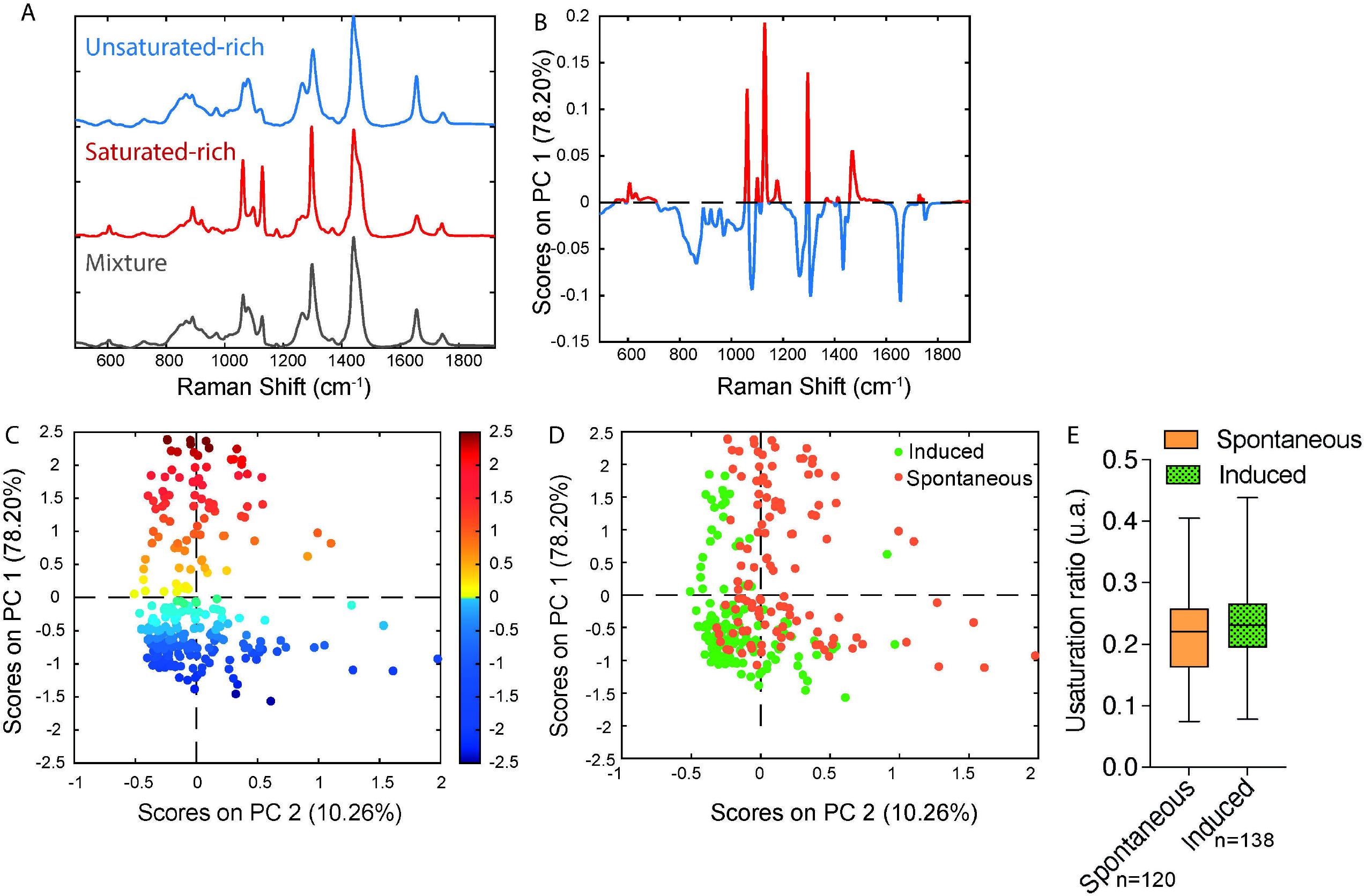
Raman microspectroscopy reveals predominance of unsaturated spectra in induced adipocytes, with heterogeneity in the unsaturation ratio for the spontaneous condition. (**A**) Representative Raman spectrum of saturated-rich, unsaturated-rich lipids and mixture identified by unguided hierarchical cluster analysis. (**B**) Loading plot PC1 indicates the discriminant peaks for saturated versus unsaturated designation, the color is associated to the attribution of specific Raman bands: saturated in blue, unsaturated in red (n=3 independent experiments, pooled for PCA analysis). (**C**) PCA score plot of Raman spectra labelled according to spontaneous (orange) and induced (green) OP9-adipocytes. (**D**) The same PCA score plot as (**C**) labelled according to the assignment of Raman spectra as saturated (red) and unsaturated (blue) lipids based on the PC1 loading determined by B. For panels C and D, each point is a single adipocyte with all its lipid droplet data points averaged. n (induced) = 138 adipocytes, 2971 spectra; n (spontaneous) =120 adipocytes, 2944 spectra. (**E**) Unsaturation ratios of spontaneous and induced at adipocyte level. The unsaturation ratio of sOP9 adipocytes is not different from iOP9 adipocytes (P=0.0855 by Mann-Whitney test).

First, we hypothesized that the smaller diameter of LDs in sOP9 adipocytes may be related to their lower unsaturation ratio, owing to the higher mobilization rate of unsaturated FAs (62). We thus plotted LD diameter versus unsaturation ratio (Figure 5A-B). The LD diameter did not correlate with the unsaturation ratio. Specifically, correlation analysis for the LD diameter to unsaturation ratio shows that the parameters are not linearly correlated in either condition (Figure 5A-B, R^2^_spont_ =0.0013; R^2^_ind_=0.0107). Even when the spectra are separated according the 3 types of lipids (saturated-rich, unsaturated-rich and mixture) the LD diameter and the unsaturation ratio are not linearly correlated. All the R^2^ are inferior to 0.05 (Figure S6). Since, no correlation was observed, the LD diameter and the unsaturation ratio were explored separately.

**Figure 5:**
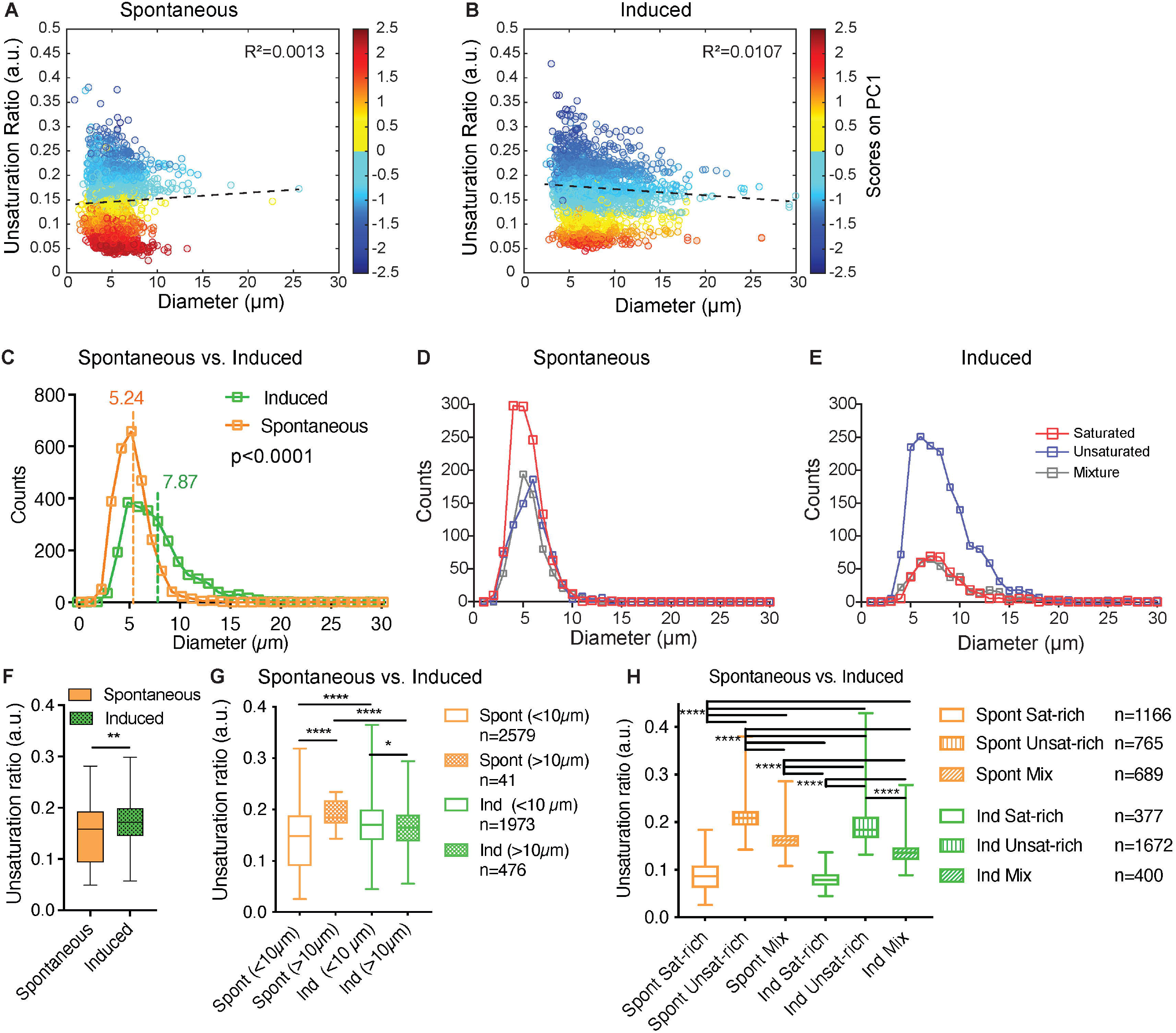
Induced adipocytes are composed of larger lipid droplets than spontaneous adipocytes, but there is no overall correlation between lipid droplet diameter and unsaturation ratio. (**A**-**B**) Unsaturation ratio as a function of the diameter of the LDs for sOP9 versus iOP9 adipocytes (n=3 independent experiments, pooled for LD analysis). The low R2 for both conditions indicates that the diameter of the LDs is not correlated with the unsaturation ratio. Dashed lines represent the correlation line. The Y axis is colored as function of the score of PC1. The circles in dark blue corresponds to LDs identified as unsaturated rich. The circles in dark red correspond to LDs identified as saturated-rich. The circles in yellow and light blue correspond to LDs identified as mixture. (**C**) Histogram for frequency of LDs as a function of the diameter for the spontaneous and induced condition. Vertical lines show the mean diameter of the LD for each condition. The mean diameter of LDs is 5.24μm (±1.85μm) and 7.87μm (±3.83μm) for sOP9 adipocytes and iOP9 adipocytes, respectively. The LDs mean diameter of sOP9-adipocytes is significantly lower compared to iOP9-adipocytes (P<0.0001, Mann-Whitney test). (**D**, **E**) When separated according to LD type a trend toward larger unsaturated lipid droplets in the induced condition is observed. (**F**) Unsaturation ratios of sOP9 versus iOP9 at LD level. LDs from iOP9 adipocytes have a higher mean unsaturation ratio compared to LDs from sOP9 adipocytes. (P<0.0014, Mann-Whitney test). (**G**) At single droplet resolution, larger droplets (>10*μ*m) have a higher unsaturation ratio as compared to smaller lipid droplets, while smaller droplets (<10*μ*m) in induced OP9-adipocytes (n=1973) have a higher unsaturation ratio than those of spontaneous OP9-adipocytes (n=2579). Highest unsaturation ratio overall is seen in large droplets from spontaneous OP9-adipocytes (n=41). (**F**) Unsaturation ratio as a function of the condition (spontaneous and induced) and the nature of the lipid (saturated-rich, unsaturated-rich and mixture). *P<0.05, ****P<0.001 by Kruskal-Wallis test. The lipid droplet of spontaneous unsaturated-rich have the higher unsaturation ratio compared to induced unsaturated-rich.

The averaged LD diameter was significantly lower (p<0.0001) in the sOP9-adipocytes (5.24±1.85*μ*m) than the iOP9-adipocytes (7.87±3.83*μ*m) (Figure 5C). When we further separated LDs by type of lipids, we found that saturated-rich, unsaturated-rich or mixture LDs are not different in their average composition when originating from iOP9 or sOP9 adipocytes, but unsaturated LDs are more frequent in iOP9s while saturated LDs are more frequent in the sOP9 condition (Figure 5D-E, Figure S5B). Unsaturated-rich LDs are overall bigger in size than saturated-rich LDs, with a predominance in iOP9 adipocytes, particularly above the diameter of 10 *μ*m. In iOP9 adipocytes, the number of unsaturated-rich LDs is higher compared to sOP9 adipocytes for LD diameter superior to 10 *μ*m.

Then, the averaged unsaturation ratio per condition were obtained from individual spectra of LDs. The unsaturation ratio of iOP9 condition (0.17±0.05) is significantly higher than sOP9 condition (0.14±0.05, p=0.0014). Also on average, LDs of iOP9 condition have more unsaturated-rich lipids compared to the LDs in sOP9 condition. Since differences are observed between sOP9 and iOP9 around a LD diameter of 10 *μ*m (Figure 5E), the unsaturation ratio of LDs was investigated as function of the threshold of 10 *μ*m (Figure 5G). The larger LDs (>10*μ*m) have the highest unsaturation ratio compared to smaller LDs. Smaller LDs (<10*μ*m) from iOP9 adipocytes have a higher unsaturation ratio compared to sOP9-adipocyte. Then, we investigated the unsaturation ratio as a function of the 3 types of LDs, namely saturated-rich, mixture or unsaturated-rich. Although we found consistent differences between the three classes of LDs, there were no significant differences between LDs of the same class in sOP9 adipocytes versus iOP9-adipocytes (Figure 5H). Not surprisingly, mixture LDs showed the greatest heterogeneity in their unsaturation ratio (Figure 5H). Overall, we conclude from this data that iOP9 adipocytes contain significantly bigger LDs and more frequently unsaturated LDs than sOP9 adipocytes. *In vivo*, both higher LD diameter and higher FA unsaturation are characteristics of cBMAds, while smaller LD diameter and lower unsaturation ratio are characteristics of rBMAds. We could not find a consistent relationship between LD diameter and unsaturation ratio.

### 3.4 Hematopoietic support capacity of sOP9-adipocytes is superior to that of iOP9-adipocytes

Another *in vivo* property of rBMAds as compared to cBMAds is their differential capacity to serve as a supportive niche to hematopoietic progenitor cells. Indeed, cBMAds are associated to the hematopoietic poor -yellow-regions of the marrow, while rBMAds are associated to the hematopoietic-rich -red-regions of the marrow. To determine whether sOP9 versus iOP9 cultures behave different in terms of hematopoietic support, we co-cultured murine hematopoietic stem and progenitor cells (HSPCs) isolated as cKit^+^Lin^-^Sca1^+^ (KLS) cells for seven days with the differentiated OP9 cells at a ratio of 1 HSPC to 10 OP9 cells. As in previous work (63), this high ratio of KLS to OP9 cells serves to avoid dispersion due to HSPC heterogeneity and avoids downstream stochastic, clone effects on analysis secondary to differences in the content of the most primitive and thus proliferative HSPCs across wells, which although small, would be exponentially exaggerated after co-culture. At the end of the seven-day co-culture period, flow cytometric analysis revealed that total hematopoietic expansion of 2,000 seeded HSPCs was 4.5-fold lower (p<0.0001) when iOP9 cultures were used as feeder as compared to the sOP9 condition (Figure 6A). Of note, total hematopoietic cells in suspension were significantly lower in the iOP9 versus sOP9 condition (p<0.05) whereas the more immature progenitor cells that are typically in close contact and adherent to the feeder cells were proportionally lower in the sOP9 condition. This suggests that iOP9 cultures may preferentially favor support of the more primitive hematopoietic progenitors, which is congruent with previous data from purified human BMAd as compared to undifferentiated stromal cells (8). The more hematopoietic progenitor supportive function of the sOP9 condition is also illustrated by the quantification of colonies formed by day 7 of co-culture (Figure 6B). The total number of hematopoietic colonies per initial number of HSPCs seeded, was significantly higher in the sOP9 coculture (0.13 colonies per initial KLS seeded) than in the iOP9 coculture (0.01 colonies per initial KLS seeded). All three colony categories, from the shiny consisting of more mature hematopoietic cells to the mixed and cobblestone colonies which contain the most immature hematopoietic progenitors were significantly increased (P<0.05) in the sOP9 condition. To understand to which extent secreted factors from either condition contributed to hematopoietic expansion, HSPCs were cultured for two days in the absence of feeder cells but in the presence of different concentrations of 48-hour conditioned media from either uOP9, sOP9 or iOP9 cultures. For the iOP9 condition we tested both a short (5+2 days) and a long (17+2 days) DMI adipogenesis induction protocol (Figure 7C). Hematopoietic expansion was greatest for HSPCs cultured in the highest concentration of undifferentiated-conditioned medium. HSPCs cultured in conditioned medium from iOP9-adipocytes did not show a significant expansion after two days in culture, independently of the length of the iOP9 DMI induction protocol. However, the conditioned medium from sOP9 cultures was substantially more supportive to HSPC maintenance than iOP9, although less than those cultured in conditioned media from undifferentiated OP9 cells. This effect increased with higher ratios of conditioned-to-basal medium. We therefore conclude that the *in vitro* hematopoietic progenitor support capacity of sOP9 cultures is superior to that of iOP9 cultures, aligning with the *in vivo* functionality of regulated versus constitutive BMAds.

**Figure 6:**
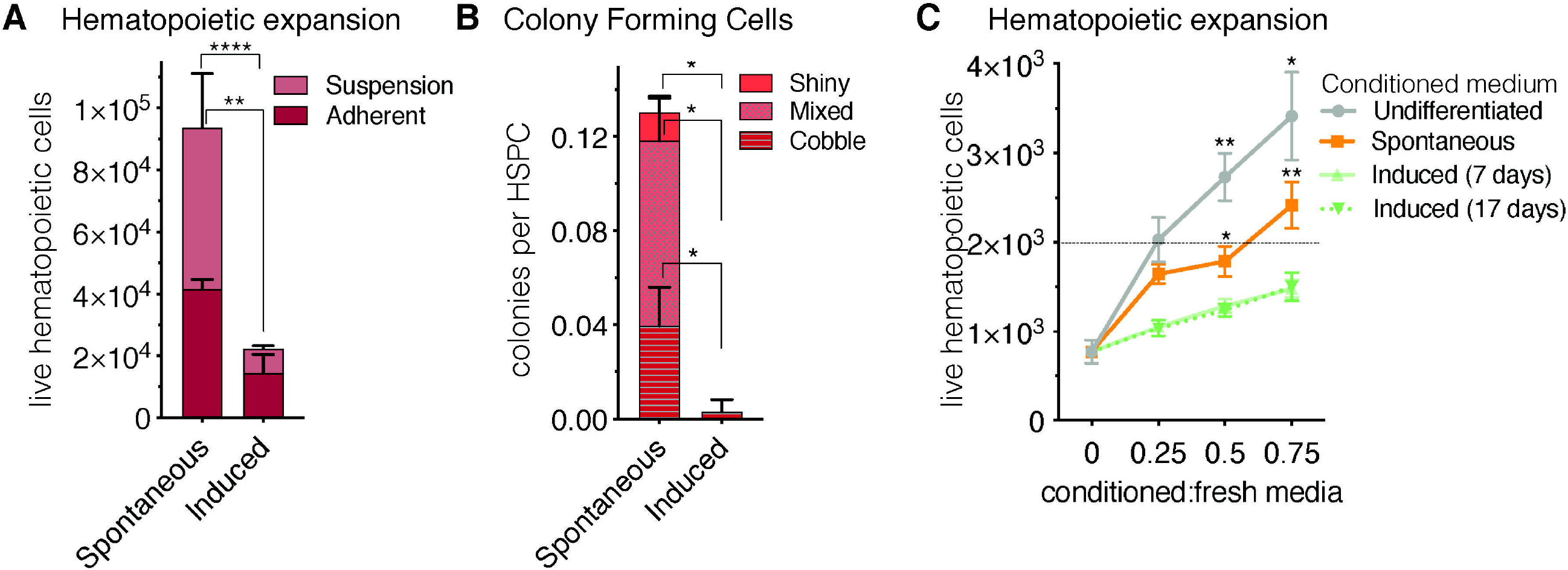
Short-term co-culture of OP9-adipocytes with hematopoietic stem and progenitor cells favours hematopoietic expansion in the spontaneous condition. (**A**) Total hematopoietic cell expansion of 2000 initial HSPCs after seven days in co-culture (including cells in adherent and cells in suspension) is significantly higher (p<0.0001) when hematopoietic stem and progenitor cells, HSPCs, (cKit^+^Lin^-^Sca1^+^, KLS) are co-seeded with spontaneous-compared to induced OP9 adipocytes. (**B**) The colony forming capacity of HSPCs seeded with spontaneous OP9 adipocytes is significantly higher (P<0.05) than when cultured with induced OP9 adipocytes. (**C**), Total hematopoietic expansion (in the absence of stroma) after two-day co-culture of 2000 HSPCs in conditioned medium from OP9 cultures comprising increasing ratios of conditioned-to-fresh media. ****P<0.0001, **P<0.01, *P<0.05 by Bonferroni’s multiple comparisons test. Error bars represent mean ± s.d. (n=3 biological replicates).

**Figure 7:**
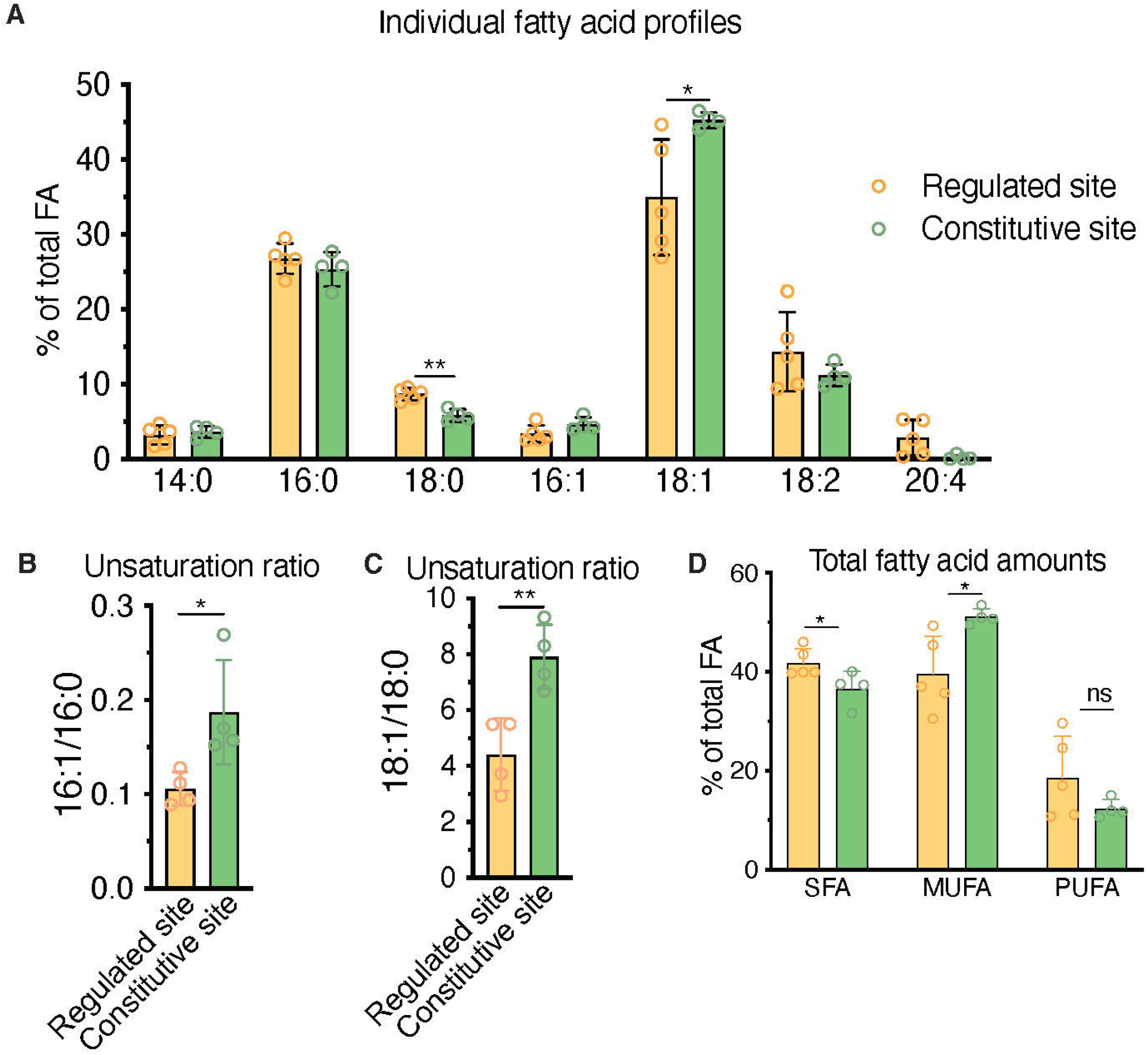
HPLC fatty acid quantification of primary human BM samples confirms a higher unsaturation ratio in constitutive BMAd-rich marrow (femoral head) versus regulated BMAd-rich marrow (ileac crest post-chemotherapy). **A-C**) HPLC analysis of lipids isolated from human iliac crest bone marrow aspirates after induction chemotherapy (regulated site, labile marrow) or from the bone marrow of femoral specimens collected upon hip replacement surgery (constitutive site, stable marrow). (**A**) Higher unsaturation content in stable versus labile marrow oil reveal nuances of individual saturated, mono-, and polyunsaturated lipid species. (**B-C**) Greater unsaturation ratios in the human marrow from the constitutive site versus the regulated siten. (**D**) **P<0.01, *P<0.05 by student’s t-test. Error bars represent mean ± s.d. (n=4 experiments from independent donors). Regulated site (iliac crest): 3 females, 1 male (age = 55 ± 12 years). Constitutive site (hip replacement): 3 males, 1 female (age = 68 ± 10 years).

### 3.5 HPLC fatty acid quantification in primary human samples confirms a higher unsaturation ratio in constitutive BMAd-rich marrow (femoral head) versus regulated BMAd-rich marrow (iliac crest post-chemotherapy)

In order to contextualize and validate the value of the FA unsaturation ratio used throughout this work as a surrogate marker for regulated versus constitutive BMAds, we set to test whether we could find differences in the unsaturation ratio of lipids extracted from one of the most extreme BM remodeling scenarios. Patients suffering from acute leukemias receive intensive aplastic chemotherapy, a regime that eliminates leukemic cells but also virtually all short-term hematopoietic progenitors in the BM, leading to a rapid adipocytic conversion of the marrow. BMAds formed in this context should represent a stereotypical example of rBMAd, and we thus set out to compare the unsaturation ratio of lipids obtained from iliac crest samples from these patients as compared to the lipid fraction obtained from surgical debris of adult patients undergoing elective hip replacement surgery, which is thought to represent a cBMAd-rich site. Analysis of these primary human BM lipid samples reflected a preferential enrichment of MUFAs in the post-chemotherapy iliac crest BM samples (regulated BMAd site) versus the femoral head BM samples (constitutive BMAd site) from hip replacement surgery (39.6% versus 51.2%, p=0.02) to the detriment of PUFAs (18.6% versus 17.0% p=0.80) and SFAs (41.8% and 36.6%, p=0.04) (Figure 7D). While rapid conversion to adipocytic marrow is induced by chemotherapy, and thus iliac crest samples are interpreted as rBMAd-rich, the differences in fatty acid profiles displayed could also be an inherent function of the anatomical site. The largest difference between FA species in the regulated versus constitutive marrow sites was seen in the 18:1 to 18:0 unsaturation ratio (4.40 versus 7.91, p=0.007), and the 16:1 versus 16:0 unsaturation ratio (0.11 versus 0.19, p=0.031) (Figure 7B-C). We thus conclude that human BM samples from a rBMAd-rich site in the context of BM remodeling driven by hematopoietic demand present a lower unsaturation ratio, as measured by 16:1/16:0 -and 18:1/18:0-FA content, than samples obtained from a cBMAd-rich site, which complements previously reported results for rabbit and rats marrow at steady-state (1,33–35) as summarized in Table 1.

Taken together, we conclude that the spontaneous versus induced OP9 *in vitro* adipogenesis model presented here replicates key differences in lipid droplet size, FA unsaturation ratio and hematopoietic support respectively associated *in vivo* to regulated versus constitutive BMAd subtypes. We additionally validate the value of the 16:1/16:0 FA unsaturation ratio as a surrogate marker to segregate rBMAd-rich versus cBMAd-rich tissues both *in vivo* and for our *in vitro* model, and the feasibility of Raman microspectroscopy to reveal equivalent differences in the unsaturation ratio at the single adipocyte level.

## 4 Discussion

BMAds constitute a distinct fat depot with low responsiveness to cold and insulin, overall relative resistance to lipolysis, and high expression of adiponectin as compared to subcutaneous fat depots (37, 65, 66). Two BMAd subtypes have been described based on their differential response to physiological and metabolic stimuli: the more stable so-called constitutive (cBMAd) depots, preferentially located in the distal parts of the skeleton, and the more labile so-called regulated (rBMAd) adipocytes, which are interspersed within the hematopoietic marrow in more proximal skeletal sites. Due to their fragility and location within the bone cavity, access to live BMAd cells for functional and cell trajectory analysis is not trivial, especially for murine BMAd depots where BM adipose tissue volume is very limiting. Although possibly limited on the extrapolation of specific metabolic behaviors (37), *in vitro* models of adipogenesis from undifferentiated stromal progenitors have been proposed as a complementary tool to study BMAd at the cellular level, so as to facilitate both gene function studies and cell behavior analysis (reviewed in (32, 67)). To our knowledge, no *in vitro* model has been described to recapitulate some or all of the differential characteristics of the *in vivo* defined rBMAd versus cBMAd depots. Here we present a model of BM adipogenesis in the form of a differential *in vitro* culture system for BM-derived OP9 stromal cells. When classically induced by an adipogenic cocktail (iOP9s) OP9-derived adipocytes resemble cBMAds and, when allowed to spontaneously differentiate by confluency in serum-containing conditions (sOP9s), they resemble rBMAds in lipid droplet size, overall lipid saturation, 16:1/16:0 FA unsaturation ratio and hematopoietic supportive properties. Indeed, we validated the value of the 16:1/16:0 FA unsaturation ratio, already described as a discriminating factor for cBMAd and rBMAd depots in rabbit and rat homeostatic tissues, in both human BM and as a marker of cBMAd-like (iOP9) versus rBMAd-like (sOP9) *in vitro* cultures. Furthermore, we have extended the reproducibility of the 16:1/16:0 FA unsaturation ratio as a discriminating factor to highly remodeled rBMAd depots in the context of patients undergoing intensive ablative chemotherapy. It is worth noting that the robustness of differences in the 16:1/16:0 FA unsaturation ratio in our model (summarized in Table 1; (1, 68), as opposed to other proposed FA pairs (e.g. 18:1/18:0), is somehow surprising and may be associated to the low mobilization rate for palmitate (16:0) and palmitoleate (16:1) (69, 70). Further studies should mechanistically address the relative contribution of *de novo* lipogenesis and FA re/uptake in lipid composition differences underlying the 16:1/16:0 FA unsaturation ratio. Indeed, at the RNA level, sOP9 adipocytes had significantly higher expression of *Lpl*, which mediates fatty acid uptake for storage, while iOP9 adipocytes had increased expression of *Atgl*, the key enzyme that initiates hydrolysis of TAGs.

After HPLC-based lipid composition analysis of OP9 cultures at the cell population level, and given the relative heterogeneity of *in vitro* adipogenic cultures (67), we implemented a novel application of Raman microspectroscopy to measure lipid unsaturation heterogeneity at the single cell level, with individual LD resolution and in a label-free manner. This approach allowed us to focus lipid composition analysis exclusively on mature adipocytes and not on the remaining unlipidated stromal component. Previous studies have shown the application of Coherent anti-Stokes Raman microspectroscopy to embryonic stem cells undergoing adipogenic differentiation, showing a measure of chemical lipid saturation composition and an increase in LD accumulation with differentiation (71, 72). While, very interestingly, individual LDs within one adipocyte appear to have an inhomogeneity in their chemical composition, this was not further analyzed. Raman microspectroscopy has allowed for the characterization of LD lipid composition in animal and *in vitro* models. Lipid droplets have been measured directly in the endothelial layer of murine aortic sections by, pointing to smaller LDs induced by TNFalpha preferentially containing more unsaturated lipids whereas larger LDs were more rich in saturated lipids (73). Another Raman-based study showed interesting differences in the pattern of distribution between saturated FA species (palmitic acid −16:0- and stearic acid −18:0), which were evenly distributed in broad areas of the cell, and unsaturated FA species (oleic acid −18:1- and linoleic acid −18:2-), which were concentrated in LDs, after exposure and uptake in macrophages (74). The effect of the high fat diet was evaluated on the lipidic composition of perivascular adipocytes in a mouse model. The size of the LDs was increased as an adaptive response to the excessive lipid income. In an *in-vitro* study, the effect of TNF-α on endothelial HMEC-1 cells during 24h was monitored by Raman microspectroscopy. The composition and the distribution of cellular lipids was modified upon exposure to TNF-α, and the increase in LD size was concomitant with the formation LDs composed of unsaturated rich lipids (75, 76). However, none of these studies provide supporting statistical analysis for an univocal association between LDs size and molecular composition. In all, we found Raman microspectroscopy to be robust for the measurement of the unsaturation ratio at the lipid droplet level. Congruently to the HPLC-derived data, we could thus confirm at the single adipocyte level the higher unsaturation ratio of iOP9 lipids as compared to the sOP9 condition. It is worth noting that the overall differences in unsaturation ratio were more pronounced when measured with HPLC analysis (1.02±0.27 for iOP9-cultures versus 0.19±0.11 sOP9-cultures, p=0.0004, Figure 2E), than with Raman (0.17±0.05 for iOP9-cultures versus 0.14±0.05 sOP9-cultures, p=0.014, Figure 5F). This difference could be related to the sample bias, as HPLC analysis was done in lipid extracts from the bulk culture while Raman microspectroscopy acquired spectra from mature adipocytes only, or to the method itself. For the HPLC analysis, the unsaturation ratio is calculated from the contribution of isolated and purified lipids. On the other hand, Raman analysis is done inside the LD where lipids are mixed. The unsaturation ratio is calculated from the contribution of the mixture of lipids which hamper the signal. Indeed, the small differences we report in the unsaturation ratio are in the range of previous Raman studies cited above, although differences are more pronounced if only the more defined fractions are considered (unsaturation-rich versus saturated-rich, Figure S5).

Diseases of lipodystrophy may provide insights into the formation of BMAds and the selective accumulation of FAs that we are struggling to understand (1, 77). Congenital generalized lipodystrophy (CGL) is a family of diseases that include a complete or partial loss of adipose depots, which reveal important insights into the consequences of genetic deficiencies involved in adipocytic maturation and function (reviewed in (77)) and may point us in the direction of the underlying nature of BMAd formation. One of the best understood CGL mouse models within the context of BMAd research are the *Ptrf^-/-^* deficient mice, which contain BMAT only in the tail vertebrae and very distal tibia with a loss of BMAds at the tibia-fibula junction and proximal thereof (1,78–80). This is accompanied by a small increase in cortical bone mineral content in adult male mice and a large increase in trabecular number and cortical bone mineral content in females (1). The differential effect on BMAT upon *PTRF* loss of function, points toward a differential regulation of LD formation in regulated versus constitutive BMAds. Alternatively, differences in unsaturated lipid ratios could be involved in the specific sensitivity of BMAds to such a mutation. More recently, diphtheria-toxin mediated ablation of Adipoq-Cre expressing cells also revealed differences in BMAd plasticity throughout the long bones in “fat-free” mice. In particular BMAds were increasingly present in the mid-diaphysis with age (81, 82). Furthermore, specific depletion of BMAds through a double conditional Osterix and Adipoq-dependent strategy caused severe reduction in rBMAds with a milder reduction in the cBMAd mass, while ATGL (*Pnpla2* gene) loss in BMAd through the same strategy caused massive rBMAd expansion in the mid diaphysis (83, 84). To what extent these discrepancies are due to divergent transcriptional programs or part of a stabilization of the adipocytic differentiation program remains to be determined. Moreover, seipin-deficient lymphoblastoid cell-lines derived from patients suffering congenital generalized lipodystrophy type 2 were shown to have an increased proportion of SFAs with a decrease in MUFAs, which are the principle products of Scd1, indicating decreased Scd1 activity. LD number and size were increased compared to control (85). Impaired Δ9-desaturase activity in nematodes also led to decreased TAG accumulation and LD size that could not be rescued completely by supplemented dietary oleate (86). Therefore, it appears that LD size is at least partially driven by the ability of Scd1 to synthesize unsaturated FAs. Although our results did not show a strong correlation of LD size with lipid unsaturation, we did observe that the largest LDs were always predominantly unsaturated. Taken together, this points to tissue-specific regulation of lipid composition independently of nutrient and metabolite availability, and possibly to sequential stages of LD (and adipocyte) maturation in the sOP9 versus the iOP9 conditions. Overall, it is tempting to suggest a continuum with smaller LD being mostly saturated, an intermediate stage of LD maturation that are of largely mixed composition, and the most mature LDs that are large and unsaturated. Follow-up studies should address this hypothesis through Raman microspectroscopy in the context of inhibition versus activation of lipolysis and reuptake with relation to lipid droplet and adipocyte size. Extrapolation to the *in vivo* scenario should be done with caution, as the primary mechanism of lipid remodeling in primary human BMAd has been shown to be independent of ATGL-mediated lipolysis (37).

Finally, we found that sOP9-adipocytes are more supportive to hematopoiesis than iOP9-adipocytes, as seen by their capacity to form hematopoietic colonies and to support hematopoietic expansion. This agrees with reports of adiponectin expressing BM pre/adipocytes -including so-called Adipo-CAR cells-being necessary for hematopoietic expansion (12, 87–88), whereas mature BMAds rather inducing HSCs quiescence or reduced proliferation (7, 8, 11, 89). Of note, depletion of recently described marrow adipocyte lineage precursors (MALPs) did not affect hematopoiesis, at least in homeostasis (90). The reduced proliferation of hematopoietic progenitors in iOP9 adipocytic cocultures was replicated in a dose-dependent manner through condition media experiments (Figure 6C), pointing to availability of soluble factors as one of the implicated mechanisms. Indeed, previous data revealed a higher expression of the hematopoietic supportive factors C-X-C Motif Chemokine Ligand 12 *(Cxcl12)* and c-Kit Ligand *(KitL)* in undifferentiated OP9 cultures versus OP9-derived adipocytes analogous to iOP9s (42), and the regulation of BM stromal cell differentiation precisely via *Cxcl12* (91). Note however that the undifferentiated OP9 condition was shown even more supportive of hematopoietic proliferation than sOP9s (Figure 6). In congruence with previous *in vivo* data (11), this observation in the OP9 system suggests that hematopoiesis is favored when mature marrow adipocytes are absent. We therefore suggest an optimum of maturation along the adipocytic differentiation axis with regard to hematopoietic support, in particular through at least limited expression of Angpt1, SCF, and Cxcl12 that are attenuated in terminally differentiated adipocytes (92). Taken together, our results indicate that, at the population level, sOP9 adipocytic cultures share similarities in terms of hematopoietic support with the labile rBMAds of the red marrow, while iOP9 adipocytic cultures impair rapid hematopoietic progenitor proliferation as described for the stable cBMAds of the yellow marrow.

The mechanisms for differential hematopoietic support *in vivo* are still poorly understood, but extrapolation from the collective findings of the field suggests that cBMAd-dense regions of the distal skeleton maintain HSCs, while rBMAd-containing red marrow would contribute to expansion of their progenitors, possibly through increased FA transfer (83). Most HSCs have low mitochondrial potential and rely mainly on anaerobic glycolysis, while downstream progenitors rely on high energy production through mitochondrial oxidative phosphorylation (93–95). Previous studies have also revealed that HSCs and leukemic cells can rely on PPAR-mediated FA oxidation (96), with activation of oxidative metabolism being a predictive factor for leukemic stem cell behavior (97). Indeed, we have shown that BM stromal cells with lower overall lipid content (sOP9 cultures) associate to increased hematopoietic expansion capacity compared with more mature BMAds that are high in lipid content (iOP9 cultures). Specific differences in FA of BMAds may contribute to the nuances of differential hematopoietic support. For example, palmitic acid was shown to have a lipotoxic effect on hematopoietic-supportive osteoblasts, and high levels are present in aged and osteoporotic bone associated with decreased hematopoiesis (98, 99). This may in part explain the overt myeloproliferation that is seen on the aged skeleton. In the current study, the stromal OP9 cell line was selected for its adipocytic differentiation potential. The inherent M-CSF deficiency may influence outcomes with regard to the hematopoietic supportive function. Previous studies have shown lymphoid and myeloid support of bone marrow adipocytes, although the extent of hematopoietic support as a function of the extent of adipocytic differentiation has not been addressed. In future studies, a comparison to marrow stromal cell lines without cytokine expression defects is warranted to provide a more representative understanding of the marrow hematopoietic and adipocytic lineages.

Future studies to understand the functional metabolic differences of rBMAd-like sOP9 versus cBMAd-like iOP9 adipocytes should focus on *in vitro* lipase- and desaturase-inhibition, complemented by the study of their role in BMAd-deficient animal models (such as the Atgl-KO or Scd1-KO) to better understand the role of rBMAd vs cBMAd FA mobilization on hematopoiesis. Altogether, our results open avenues for future work on the OP9-adipocyte system to understand the mechanisms underlying BMAd heterogeneity and its impact on both hematopoiesis and bone health.

## 5 Conflict of Interest

The authors declare no conflict of interest at this time.

The authors declare that the research was conducted in the absence of any commercial or financial relationships that could be construed as a potential conflict of interest.

## 6 Author Contributions

J.T. planned experiments, analyzed and interpreted the results. G.F. performed Raman acquisitions, interpreted and analyzed the results and compiled the corresponding figures. A.D. and N.B. performed HPLC analyses. J.T., C.B. and N.D.T. performed *in vitro* culture experiments. V.C. performed co-cultures. G.F. and L.D. performed statistical analyses for Raman data. J.T., G.F., A.D., C.C. and O.N. interpreted the Raman and HPLC data. G.Q.D. and O.N. designed and optimized OP9 differentiation protocols and associated hematopoietic support assays. O.N. and G.P. initiated the project, while C.C. and O.N. co-supervised the project. O.N. and J.T. wrote the manuscript, G.F. and C.C. significantly edited the manuscript.

## 7 Funding

J.T., V.C. and O.N. were financed by Swiss National Science Foundation (SNSF) grants PP00P3_176990 and PP00P3_183725 (to O.N), the Anna Fuller cancer fund (J.T. and O.N.) and UNIL unrestricted funds to O.N.. D.N.T. was funded by the Whitaker Fellowship program in Bioengineering. C.B. was financed by Sinergia funding CRSII5_186271.

## 8 Acknowledgments

OP9 cells were generously provided by T. Nakano, Kyoto University, Japan, via the Daley Lab. MS5 stromal cells were generously provided by Prof. Gary Gilliland, C3H10T1/2 and MC3T3 stromal cells by Prof. Peter Hauschka (both at Children’s Hospital Boston), and AFT024 as well as BFC012 from Prof. Rudolf Jaenisch (Whitehead Institute, Massachusetts Institute of Technology). We are very grateful to Aurélien Oggier and Silvia Vaz Ferreira Lopes for technical help with qPCR experiments and to the support of the EPFL Flow Cytometry Core Facility for flow cytometric sorting and EPFL Bioscreening Facility for Digital Holographic Microsocopy imaging. This collaborative work was made possible by the networking opportunities facilitated by the Bone Adiposity in HEalth and Disease (BoneAHEAD) Network consortium, and by the International Bone Marrow Adiposity Society (BMAS).

## 9 Data Availability

Original datasets are available in Mendeley Data publicly accessible repository: [http://dx.doi.org/10.17632/wdm9gvz3bm.1].

## 11 Supplementary Material

Supplementary Figures are available as an annex pdf document.

## Notes

http://dx.doi.org/10.17632/wdm9gvz3bm.1

### Competing Interest Statement

The authors have declared no competing interest.

