## Supplemental Figures for "Raman microspectroscopy reveals unsaturation heterogeneity at the lipid droplet level and validates an *in vitro* model of bone marrow adipocyte subtypes"

### Supplementary Figures:

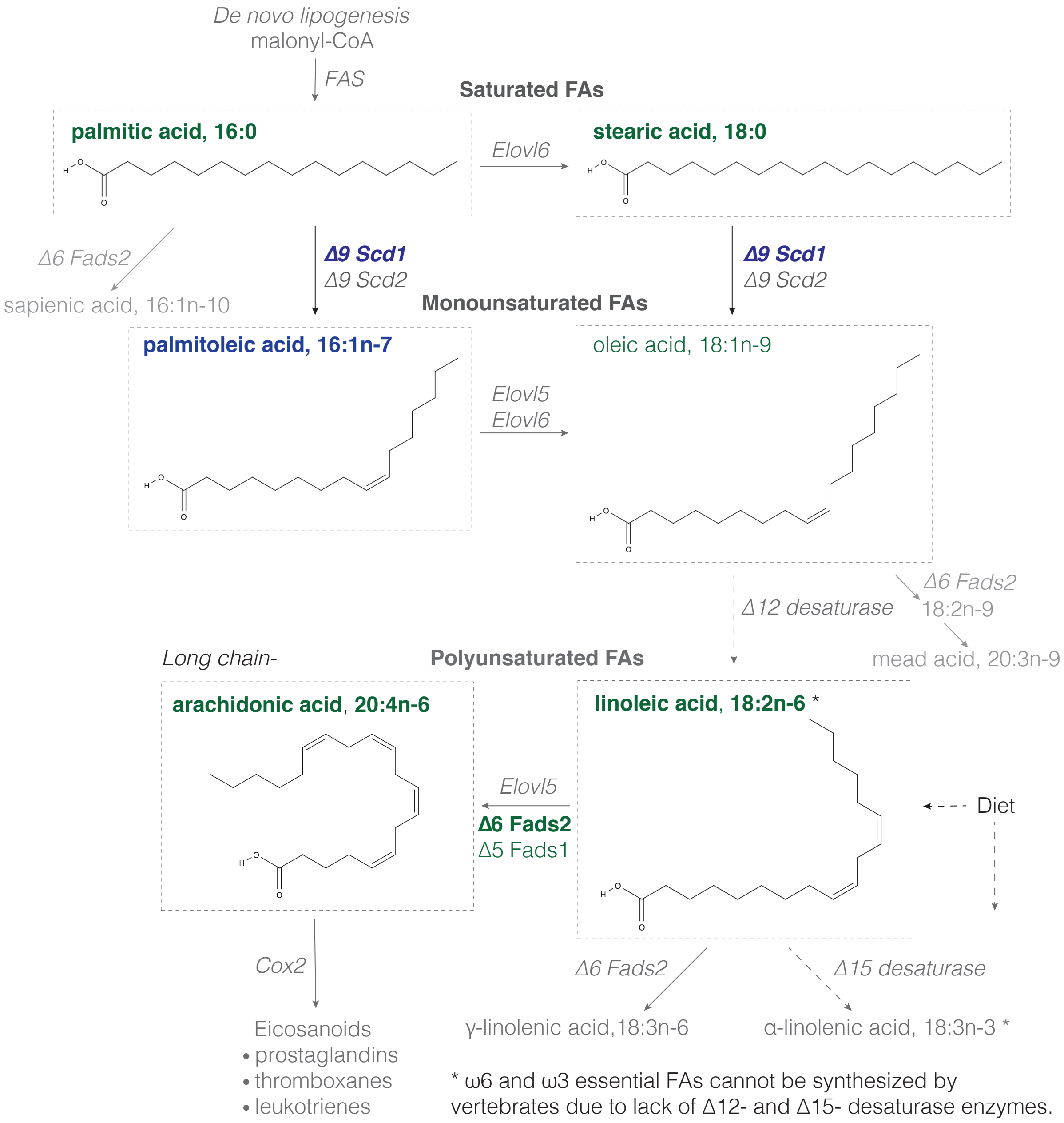


**Fig S1: Schematic of fatty acid synthesis for major lipid species in OP9 culture model.** Green: relative abundance found in spontaneous OP9 adipocytes (sOP9) by HPLC (lipid) or RT-qPCR (enzyme RNA transcript) quantification; blue: relative abundance found in induced OP9-adipocytes (iOP9) by HPLC or RT-qPCR; grey: no differenced measured by HPLC or RT-qPCR; dotted grey arrow: not a process in vertebrates. Bold green or bold blue indicates statistically significant differences and non-bold green or blue indicates quasi-significant differences. Chemical structures were generated with MolView v2.4.

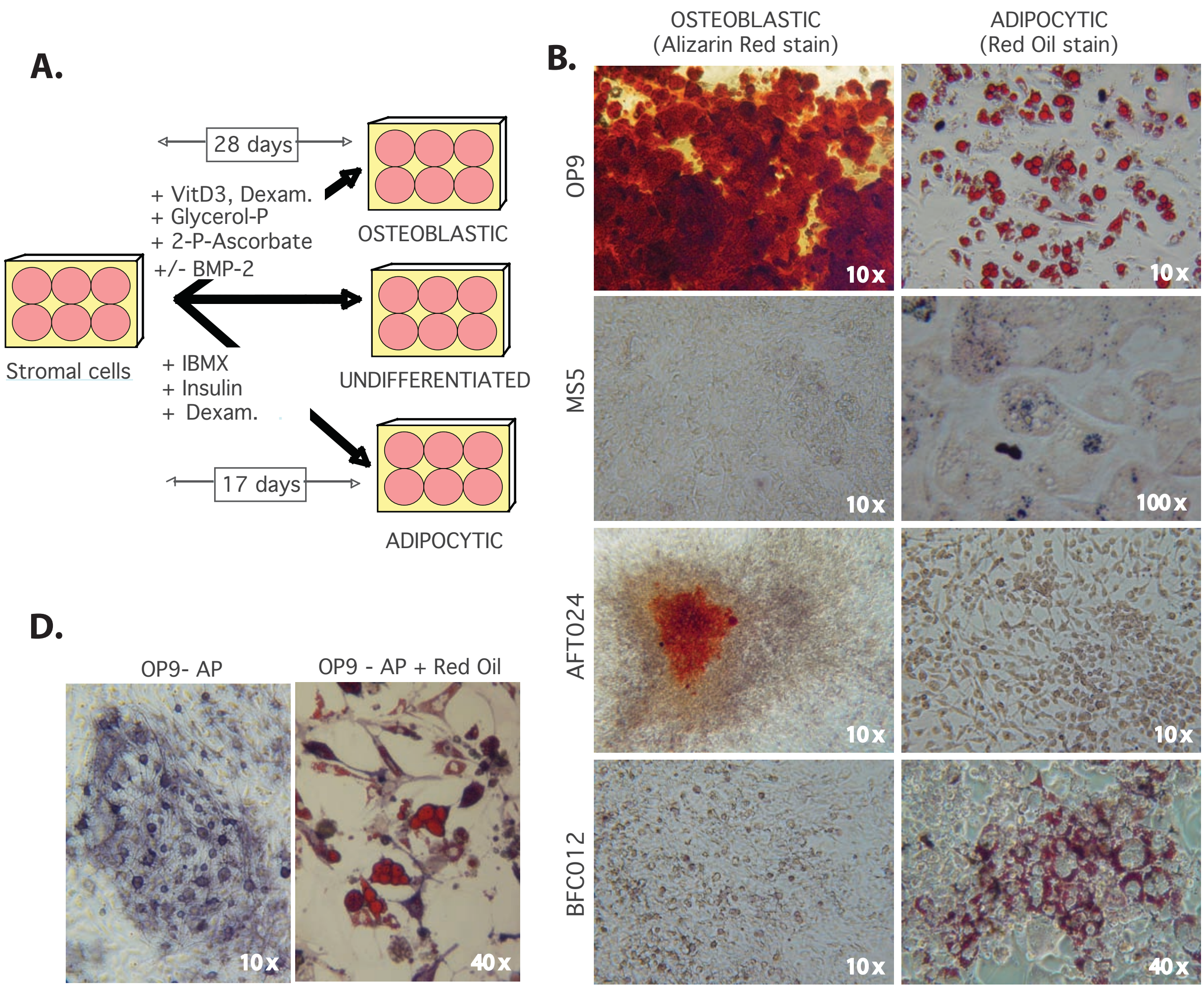


C.

| Stromal differentiation: | | Origin | Supportive | OSTEO (Alizarin Red) | ADIPO (Red Oil) |
|---|---|---|---|---|---|
| **OP9** | (n=3) | BM | √ | ++ | +++ |
| **MS5** | (n=1) | BM | √ | - | +/- |
| **C3H10T1/2** | (n=1) | BM | √ | +/- | - |
| **AFT024** | (n=2) | FL | √ | ++ | + |
| **BFC012** | (n=2) | FL | X | +/- | +/- |
| **MC3T3-E1-4** | (n=3) | BM-Osteo | ? (+/-) | ++ | - |
| **MC3T3-E1-24** | (n=1) | BM-Fibro | X | +/- | +/- |
| **Day 6 EB** | (n=1) | ES cells | N/A | +++ | - |

**Fig S2: The OP9 BM-derived stromal cell line displays optimal adipocytic differentiation with conserved osteogenic and hematopoietic-supportive function as compared to other murine stromal cell lines. (A)** Experimental approach used to induce dual osteogenic or adipocytic differentiation of murine stromal lines. **(B)** Representative micrographs of OP9, MS5, AFT024 and BFC012 stromal cell lines upon differentiation as described in (B). Osteogenic potential is revealed by Alizarin red stains for calcium deposition (left panels), and adipogenic potential is revealed by O Red Oil stain for neutral lipid accumulation (right panels). Note that O Red Oil stains of MS5 and BFC012 cells are shown at a higher magnification to appreciate the smaller lipid droplets. **(C)** Summary of the efficiency of differentiation of all cell lines tested. 'Supportive' stands for demonstrated ability to support hematopoiesis *in vitro* as revealed by formation of cobblestone colonies when co-cultured with hematopoietic stem and progenitor cells (cKit$^{+}$Lin$^{-}$Sca1$^{+}$, KLS). Semiquantitative assessment of differentiation: ++ (70-90%), ++ (40-60%), + (10-30%), +/- (<5%), - (not detected); (n=2-4 independent experiments per cell line). OP9s differentiated best into both osteoblastic and adipocytic fates while capable of efficient hematopoietic support, and were chosen for the remaining studies for this reason. **(D)** Left panel: OP9 cells were assayed after 2-week confluency for alkaline phosphatase enzymatic activity, a known marker of adventitial reticular cells (AP, purple precipitate), and reveal a delicate reticular network stain with intensified signal in morphologically mature adipocytes. Right panel: when OP9 cultures exposed to the adipocytic culture were stained for both AP enzymatic activity and lipid accumulation through O Red Oil (right panel), positivity of AP signal was confirmed in O Red Oil positive, fully lipidated mature adipocytes (see purple signal in cytoplasm around lipid droplets).

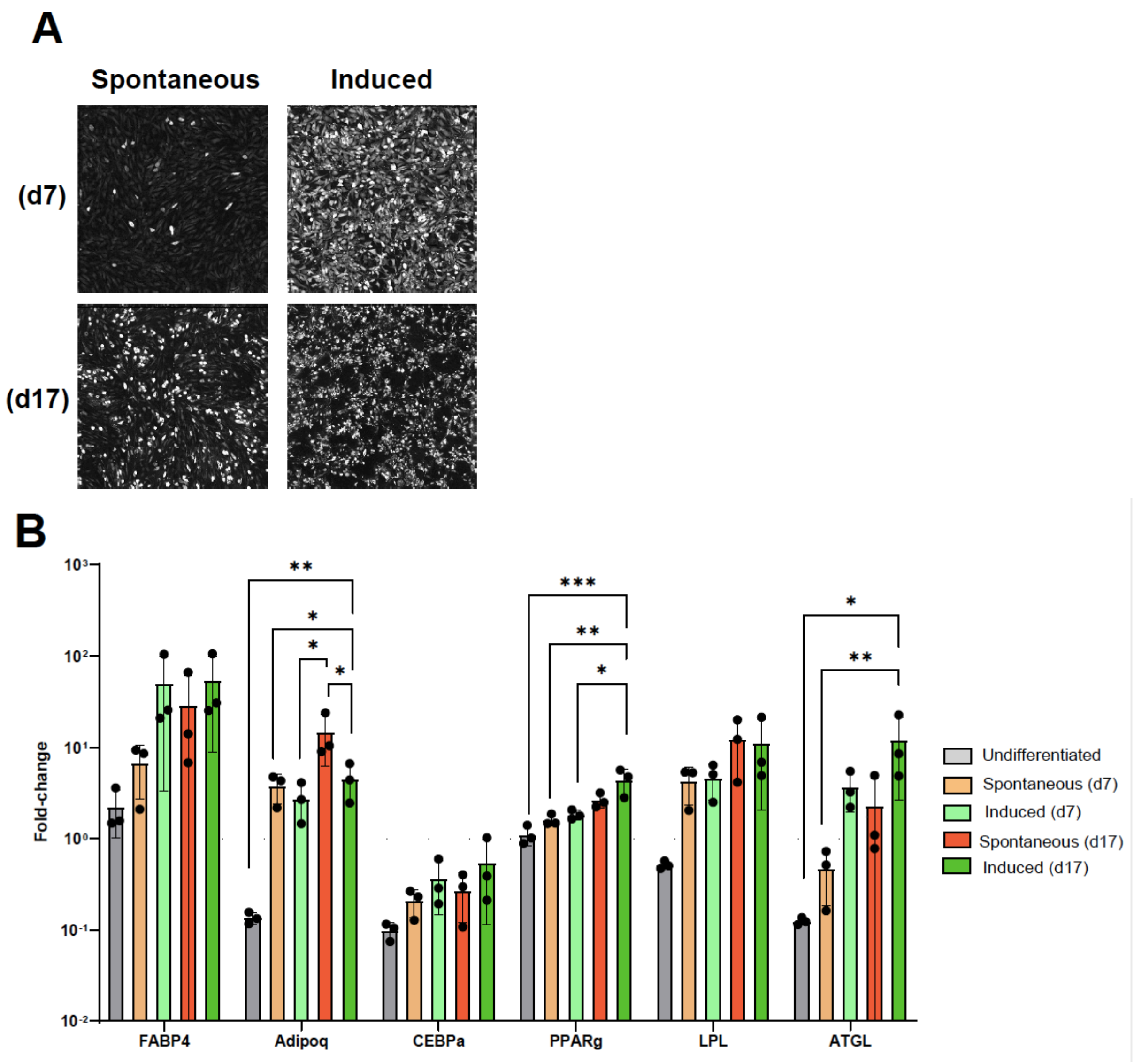


**Fig S3: Canonical adipocytic transcriptional signature for spontaneous versus induced cultures in OP9-EGFP cells. (A)** Representative digital holographic microscopy (DHM) images of OP9-EGFP cells undergoing adipocytic differentiation. Lipid accumulation is enhances in induced OP9-EGFP cells as compared to the spontaneous condition, although differentiation on Day 7 is slower than for the source OP9 cell line (see Figure 2). **(B)** RT-qPCR analysis: canonical markers of adipogenesis are increased in differentiating OP9-EGFP cells, with no significant differences at the transcriptional level between spontaneous (sOP9) and induced (iOP9) conditions at the same timepoint except for Adipoq at d17. ***P<0.001, **P<0.01, *P<0.05 by ANOVA test with Bonferroni's multiple comparisons adjustment. Error bars represent mean ± s.d. (n = 3 independent experiments)

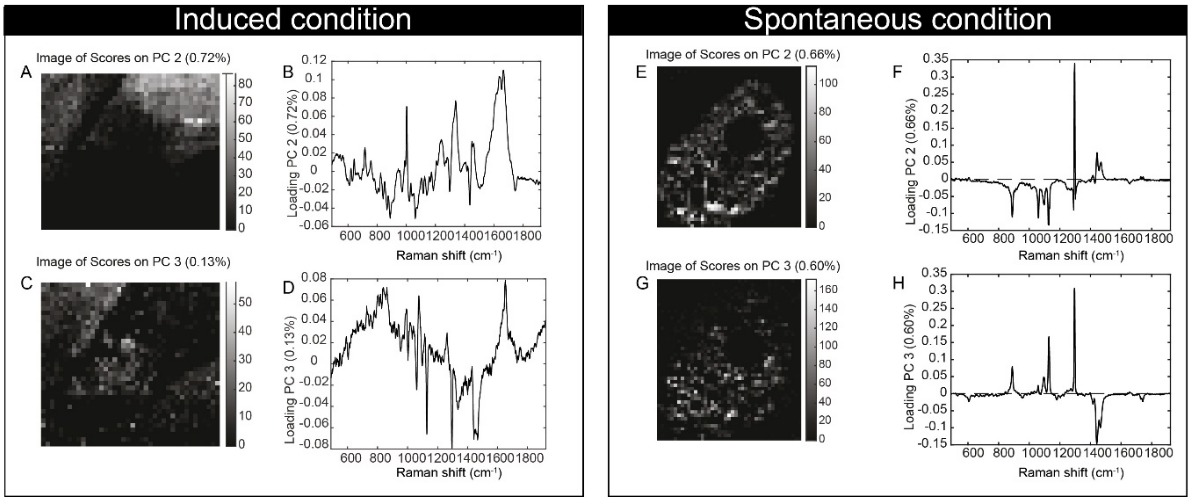


**Fig S4: Image scores and loading plots on PC2 and PC3 from Figure 3.** Induced condition: (**A)** and (**C**) Raman images of the scores of PC2 and PC3 respectively. Pixels in black represent a score of the PC equal to zero, which correspond to the absence of the spectra that drive the PC. Pixels in shades of white correspond to the contribution of PC. (**B)** and (**D**) Loadings PC2 and PC3. Spontaneous condition: **(E)** and (**G**) Raman images of the scores of PC2 and PC3 respectively. The color code is the same used in the left panel. (**F**) and (**H**) Loadings PC2 and PC3.

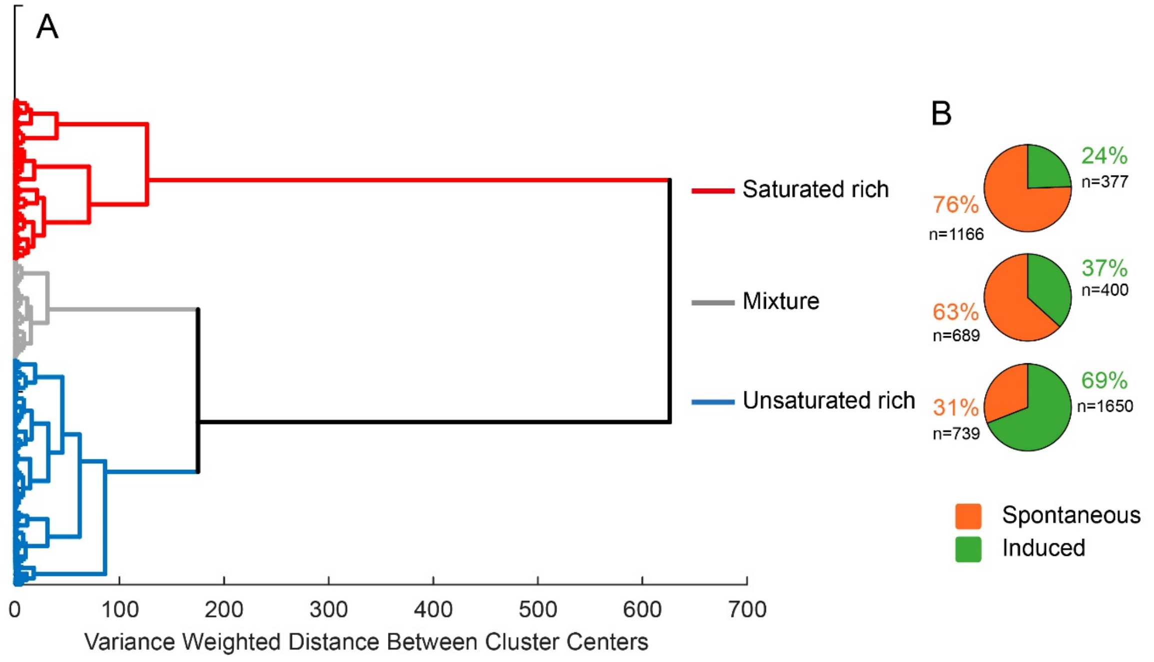


**Figure S5: Dendrogram for spectral classification.** (**A**) Raman spectra of each individual lipid droplet were classified into 3 categories: saturated-rich (red), unsaturated-rich (blue) and mixture (grey) using hierarchical cluster analysis. The dendrogram shows the classification of each spectrum into the 3 categories. (**B**) The pie charts represent the percentage of spectra per condition within each category. In the saturated-rich category, 76% of the spectra belong to the spontaneous condition and 24% belong to the induced condition. In the unsaturated-rich category, 31% of the spectra belong to the spontaneous condition and 69% belong to the induced condition. In the mixture category, 63% of the spectra belong to the spontaneous condition and 37% belong to the induced condition. The "n" corresponds to the number of lipid droplets. Due to the high number of spectra (~5000) the labels of each spectrum could not be displayed.

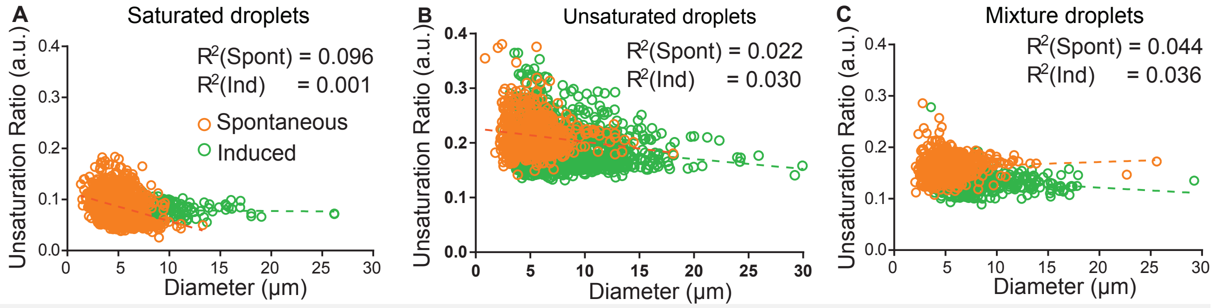


**Figure S6. Differences in lipid droplet diameter do not correspond with difference in mean unsaturation ratio in induced (green) versus spontaneous (orange) OP9-adipocytes**. (**A**) Saturated-rich droplets, (**B**) unsaturated-rich droplets, and (**C**) mixture droplets in spontaneous versus induced OP9-adipocytes show no correlation with lipid droplet diameter. The dashed lines represent the correlation lines.

#### **Supplementary Tables:**

**Table S1:** Lipidomics LC-HRMS standards and solvents

| Standard/Solvent | Supplier |
|---|---|
| Chloroform stabilized by 0.5-1% ethanol | Merck (Darmstadt, Germany) |
| Methanol (UPLC grade) | Merck (Darmstadt, Germany) |
| Formic acid (98-100%, for analysis) | Merck (Darmstadt, Germany) |
| Acetonitrile (UPLC grade) | Biosolve (Valkenswaard, Netherlands) |
| Deionized water | B. Braun (Melsungen, Germany) |
| Ammonium formate | Avanti Polar Lipids (Alabaster, AL, USA) |
| 1,2-didodecanoyl-sn-glycero-3-phosphocholine | Avanti Polar Lipids (Alabaster, AL, USA) |
| 1-heptadecanoyl-2-myristoleoyl-sn-glycero-3-phosphocholine | Avanti Polar Lipids (Alabaster, AL, USA) |
| 1-heptadecanoyl-2-myristoleoyl-sn-glycero-3-phospho-(1'-myo-inositol) | Avanti Polar Lipids (Alabaster, AL, USA) |
| 1-heptadecanoyl-2-myristoleoyl-sn-glycero-3-phospho-L-serine | Avanti Polar Lipids (Alabaster, AL, USA) |
| N-(dodecanoyl)-sphing-4-enine-1-phosphocholine | Avanti Polar Lipids (Alabaster, AL, USA) |
| N-(heptadecanoyl)-sphing-4-enine | Avanti Polar Lipids (Alabaster, AL, USA) |
| D-glucosyl-ß-1,1'N-octanoyl-D-erythro-sphingosine | Avanti Polar Lipids (Alabaster, AL, USA) |
| sn-(3-tetradecanoyl-2-hydroxy)-glycerol-1-phospho-sn-3'-(1'-tetradecanoyl-2'-hydroxy)-glycerol | Avanti Polar Lipids (Alabaster, AL, USA) |
| Bis-[2-(9Z-octadecenoyl)-3-lyso-sn-glycero]-1-phosphate | Avanti Polar Lipids (Alabaster, AL, USA) |
| 1,2-ditetradecanoyl-sn-glycero-3-phospho-(1'-sn-glycerol) | Avanti Polar Lipids (Alabaster, AL, USA) |
| 1,2-di-(9E-octadecenoyl)-sn-glycero-3-phospho-(1'-sn-glycerol) | Sigma (St. Louis, MO, USA) |
| 1,2,3-Trioctanoyl-sn-glycerol | Sigma (St. Louis, MO, USA) |
| 1,2,3-Tridecanoyl-sn-glycerol | Sigma (St. Louis, MO, USA) |
| 1,2,3-Tridodecanoyl-sn-glycerol | Sigma (St. Louis, MO, USA) |
| 1,2,3-Tritetradecanoyl-sn-glycerol | Sigma (St. Louis, MO, USA) |
| 1,2,3-Trihexadecanoyl-sn-glycerol | Sigma (St. Louis, MO, USA) |
| 1,3-Dioctadecanoyl-sn-glycerol | |
| (18:1(9Z)/18:1(9Z)/0:0)1,2-di-(9Z-octadecenoyl)-sn-glycerol | Sigma (St. Louis, MO, USA) |
| 1,2-Dihexadecanoyl-sn-glycerol | Sigma (St. Louis, MO, USA) |

**Table S2:** RT-qPCR primer sequences

| Gene | Forward primer (5' – 3') | Reverse primer (5' – 3' |
|---|---|---|
| AdipoQ | TGT TCC TCT TAA TCC TGC CCA | CCA ACC TGC ACA AGT TCC CTT |
| Atgl | TGA CCA TCT GCC TTC CAG A | TGT AGG TGG CGC AAG ACA |
| C/ebpa | GGT GGA CAA GAA CAG CAA CGA | GCG GTC ATT GTC ACT GGT CA |
| Elovl1 | CAT GCT TTC CAA GGT CAT TGA GCT G | TCT CAG TTG GCC TGACC TTGGTGG |
| Fads1 | CCA GAT TGA ACA CCA CCT CTT | GAC TCA TAC TTG ATG CCG TAC TT |
| Fads2 | CAC AAG GAC CCG GAC ATA AA | TGG TTG TAG GGC AGG TAT TTC |
| Fabp4 | ATGTGCGACCAGTTTGTG | TTT GCC ATC CCA CTT CTG |
| Lpl | GGC CGC CCT GTA CAA GAG A | AAC TCC TCC TCC ATC CAG TTG A |
| Pparg | CCT GCA TCT CCA CCT TAT TAT TCT | AAA CCC TTG CAT CCT TCA CA |
| Rpl13 | CTC ATC CTG TTC CCC AGG AA | GGG TGG CCA GCT TAA GTT CTT |
| Scd1 | TTC CCT CCT GCA AGC TCT AC | CAG AGC GCT GGT CAT GTA GT |
| Scd2 | GGC CCA CAT ACT GCA AGA G | TTC AAA CTT CTC GCC TCC AT |
